# Cross-Protection Induced by Highly Conserved Human B, CD4^+,^ and CD8^+^ T Cell Epitopes-Based Coronavirus Vaccine Against Severe Infection, Disease, and Death Caused by Multiple SARS-CoV-2 Variants of Concern

**DOI:** 10.1101/2023.05.24.541850

**Authors:** Swayam Prakash, Nisha R. Dhanushkodi, Latifa Zayou, Izabela Coimbra Ibraim, Afshana Quadiri, Pierre Gregoire Coulon, Delia F Tifrea, Berfin Suzler, Mohamed Amin, Amruth Chilukuri, Robert A Edwards, Hawa Vahed, Anthony B Nesburn, Baruch D Kuppermann, Jeffrey B. Ulmer, Daniel Gil, Trevor M. Jones, Lbachir BenMohamed

## Abstract

**Background:** The Coronavirus disease 2019 (COVID-19) pandemic has created one of the largest global health crises in almost a century. Although the current rate of SARS-CoV-2 infections has decreased significantly; the long-term outlook of COVID-19 remains a serious cause of high death worldwide; with the mortality rate still surpassing even the worst mortality rates recorded for the influenza viruses. The continuous emergence of SARS-CoV-2 variants of concern (VOCs), including multiple heavily mutated Omicron sub-variants, have prolonged the COVID-19 pandemic and outlines the urgent need for a next-generation vaccine that will protect from multiple SARS-CoV-2 VOCs.

**Methods:** In the present study, we designed a multi-epitope-based Coronavirus vaccine that incorporated B, CD4^+^, and CD8^+^ T cell epitopes conserved among all known SARS-CoV-2 VOCs and selectively recognized by CD8^+^ and CD4^+^ T-cells from asymptomatic COVID-19 patients irrespective of VOC infection. The safety, immunogenicity, and cross-protective immunity of this pan-Coronavirus vaccine were studied against six VOCs using an innovative triple transgenic h-ACE-2-HLA-A2/DR mouse model.

**Results:** The Pan-Coronavirus vaccine: (*i*) is safe; (*ii*) induces high frequencies of lung-resident functional CD8^+^ and CD4^+^ T_EM_ and T_RM_ cells; and (*iii*) provides robust protection against virus replication and COVID-19-related lung pathology and death caused by six SARS-CoV-2 VOCs: Alpha (B.1.1.7), Beta (B.1.351), Gamma or P1 (B.1.1.28.1), Delta (lineage B.1.617.2) and Omicron (B.1.1.529). <u>Conclusions</u>: A multi-epitope pan-Coronavirus vaccine bearing conserved human B and T cell epitopes from structural and non-structural SARS-CoV-2 antigens induced cross-protective immunity that cleared the virus, and reduced COVID-19-related lung pathology and death caused by multiple SARS-CoV-2 VOCs.

## INTRODUCTION

While the Wuhan Hu1 variant of SARS-CoV-2 is the ancestral reference virus, Alpha (B.1.1.7), Beta (B.1.351), Gamma or P1 (B.1.1.28.1), and Delta (lineage B.1.617.2) variants of concern (VOCs) subsequently emerged in Brazil, India, and South Africa vaccines from 2020 to 2022 (1). The most recent SARS CoV-2 variants, including multiple heavily mutated Omicron (B.1.1.529) sub-variants, have prolonged the COVID-19 pandemic (2–6). These new variants emerged since December 2020 at a much higher rate, consistent with the accumulation of two mutations per month, and strong selective pressure on the immunologically important SARS-CoV-2 genes (7). The Alpha, Beta, Gamma, Delta, and Omicron Variants are defined as Variants of Concern (VOC) based on their high transmissibility associated with increased hospitalizations and deaths (8). This is a result of reduced neutralization by antibodies generated by previous variants and/or by the first-generation COVID-19 vaccines, together with failures of treatments and diagnostics (9, 10). Dr. Peter Marks, Director/CBER (Center for Biologics Evaluation and Research) for the FDA recently outlined the need for a next-generation vaccine that will protect from multiple SARS-CoV-2 VOCs (11, 12).

Besides SARS CoV-2 variants, two additional Coronaviruses from the severe acute respiratory syndrome (SARS) like betacoronavirus (sarbecovirus) lineage, SARS coronavirus (SARS-CoV) and MERS-CoV, have caused epidemics and pandemics in humans over the past 20 years (13). In addition, the discovery of diverse Sarbecoviruses in bats together with the constant “jumping” of these zoonotic viruses from bats to intermediate animals raises the possibility of another COVID pandemic in the future (14–19). Hence, there is an urgent need to develop a pre-emptive universal pan-Coronavirus vaccine to protect against all SARS-CoV-2 variants, SARS-CoV, MERS-CoV, and other zoonotic Sarbecoviruses with the potential to jump from animals into humans.

The Spike protein is a surface predominant antigen of SARS-CoV-2 that is involved in the docking and penetration of the virus into the target host cells (20–22). As such, the Spike protein is the main target of the first-generation COVID-19 subunit vaccines aiming mainly at inducing neutralizing antibodies (23, 24). Nearly 56% of the 10 billion doses of first-generation COVID-19 vaccines are based on the Spike antigen alone(25), while the remaining 44% of the COVID-19 vaccines were based on whole virion inactivated (WVI) vaccines (26, 27). Both the Spike-based COVID-19 sub-unit vaccines and the whole virion-inactivated vaccines were successful (20–22). However, because the Spike protein is the most mutated SARS-CoV-2 antigen, these first-generation vaccines lead to immune evasion by many new variants and subvariants, such as Omicron XBB1.5 sub-variant (25), (28, 29). Therefore, the second-generation COVID-19 vaccines should be focused not only on the highly variable Spike protein but also on other highly conserved structural and non-structural SARS-CoV-2 antigens capable of inducing protection mediated by not only neutralizing antibodies but also by cross-reactive CD4^+^ and CD8^+^ T cells (30–33).

We have previously mapped and characterized the antigenicity and immunogenicity of genome-wide B cell, CD4^+^ T cell, and CD8^+^ T cell epitopes that are highly conserved and present a larger global population coverage (33). We hypothesize that multi-epitope vaccine candidates that express these highly conserved, antigenic, and immunogenic B and T cell epitopes will protect against multiple SARS-CoV-2 VOCs. The present study: (1) Identified seven B cell epitopes, six CD4^+^ T cell epitopes, and sixteen CD8^+^ T cell epitopes that are highly conserved within (*i*) 8.7 million genome sequences of SARS-CoV-2, (*ii*) all previous and current SARS-CoV-2 Variants; (*iii*) SARS-CoV; (*iv*) MERS-CoV; (*v*) common cold Coronaviruses (HKU, OC1,); and (*vi*) in animal CoV (i.e., Bats, Civet Cats, Pangolin and Camels); (2) Established that those epitopes were selectively recognized by B cells, CD4^+^ T cells, and CD8^+^ T cells from “naturally protected” asymptomatic COVID-19 patients; and (3) Demonstrated that a multi-epitope pan-Coronavirus vaccine that includes the above B cell, CD4^+^ T cell, and CD8^+^ T cell epitopes generated cross-protection against all the five known SARS-CoV-2 VOCs i.e., SARS-CoV-2 (USA-WA1/2020), Alpha (B.1.1.7), Beta (B.1.351), Gamma (P.1), Delta (B.1.617.2), and Omicron (B.1.1.529) in a novel triple transgenic HLA-A*02:01/HLA-DR hACE-2 mouse model of COVID-19.

## RESULTS

### 1. Highly conserved SARS-CoV-2 epitopes are selectively recognized by CD8^+^ and CD4^+^ T-cells from asymptomatic COVID-19 patients irrespective of variants of concern infection

To identify “universal” SARS-CoV-2 epitopes to be included in a multi-epitope pan-Coronavirus Vaccine; we previously screened the degree of conservancy for human CD8^+^ T cell, CD4^+^ T cell, and B-cell epitopes that span the whole SARS-CoV-2 genome (33). CD8^+^ T cell epitopes were screened for their conservancy against variants namely h-CoV-2/Wuhan (MN908947.3), h-CoV-2/WA/USA2020 (OQ294668.1), h-CoV-2/Alpha(B1.1.7) (OL689430.1), h-CoV-2/Beta(B 1.351) (MZ314998), h-CoV-2/Gamma(P.1) (MZ427312.1), h-CoV-2/Delta(B.1.617.2) (OK091006.1), and h-CoV-2/Omicron(B.1.1.529) (OM570283.1) (33). We observed 100% conservancy in all the SARS-CoV-2 variants of concern for 14 of our 16 predicted CD8^+^ T cell epitopes (ORF1ab_2210-2218_, ORF1ab_3013-3021_, ORF1ab_4283-4291_, ORF1ab_6749-6757_, ORF6_3-11_, ORF7b_26-34_, ORF10_3-11_, ORF10_5-13_, S_958-966_, S_1000-1008_, S_1220-1228_, E_20-28_, M_52-60_, and M_89-97_) (**Fig. S1**) and (33). The only exceptions were epitopes E_26-34_ and ORF8a_73-81_ which showed an 88.8% conservancy against Beta (B.1.351) and Alpha (B.1.1.7) variants respectively (**Fig. S1**) and (33). All of the 6 highly immunodominant “universal” CD4^+^ T cell epitopes (ORF1a_1350-1365_, ORF6_12-26_, ORF8b_1-15_, S_1-13_, M_176-190_, and N_388-403_), we previously reported (33), remained 100% conserved across all the SARS-CoV-2 VOCs (**Fig. S2**).

Next, we determined whether the highly conserved “universal” CD8^+^ and CD4^+^ T cell epitopes were differentially recognized by T cells from asymptomatic (ASYMP) versus symptomatic (SYMP) COVID-19 patients. We compared the magnitude of CD8^+^ and CD4^+^ T cell responses specific to each of the conserved epitopes among 38 ASYMP and 172 SYMP COVID-19 patients. We recruited COVID-19 patients infected with SARS-CoV-2 Beta (B.1.351) and SARS-CoV-2 Omicron (B.1.1.529) spanning two years of the COVID-19 pandemic (**Fig. 1A**). Fresh PBMCs were isolated from SYMP and ASYMP COVID-19 patients, on average within 4 days after reporting their first symptoms. PBMCs were then stimulated *in vitro* for 72 hours using each of the 16 CD8^+^ T cell epitopes or each of the 6 CD4^+^ T cell epitopes. Numbers of responding IFN-γ-producing CD8^+^ and CD4^+^ T cells (quantified in ELISpot assay as the number of IFN-γ-spot forming cells, or “SFCs”) were subsequently determined.

**Figure 1.**
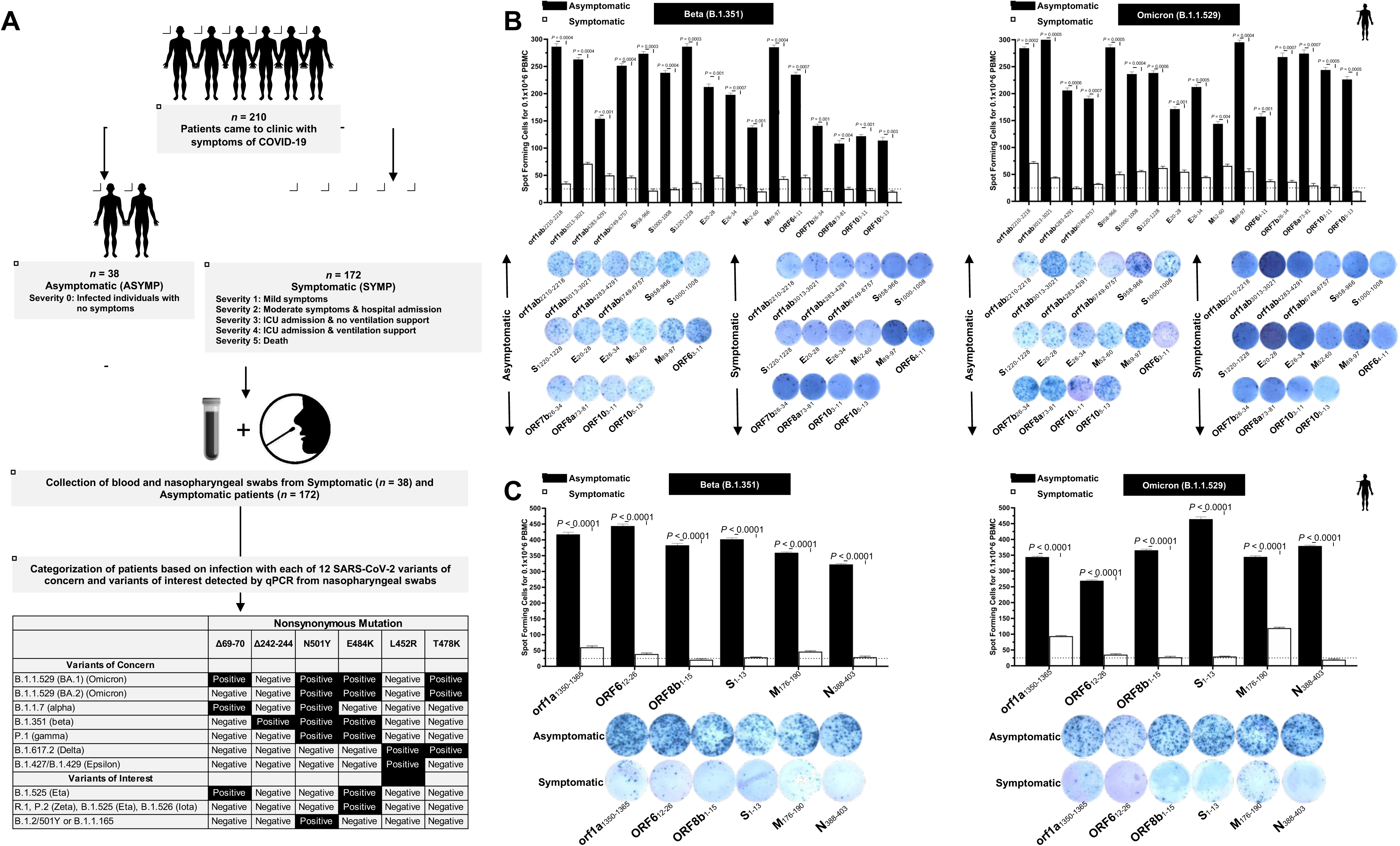
Screening of COVID-19 patients based on SARS-CoV-2 variants and subsequent evaluation of IFN-γ CD8^+^ and CD4^+^ T cell responses for conserved CD8^+^, and CD4^+^ T cell “asymptomatic” epitopes: (**A**) Experimental plan showing screening process of COVID-19 patients (*n* = 210) into Asymptomatic and Symptomatic categories based on clinical parameters. Blood and nasopharyngeal swabs were collected from all the subjects and a qRT-PCR assay was performed. Six novel nonsynonymous mutations (Δ69-70, Δ242-244, N501Y, E484K, L452R, and T478K) were used to identify the haplotypes unique to different SARS-CoV-2 variants of concern (Omicron (B.1.1.529 (BA.1)), Omicron (B.1.1.529 (BA.2)), Alpha (B.1.1.7), Beta (B.1.351), Gamma (P.1), Delta (B.1.617.2), and Epsilon (B.1.427/B.1.429)) and variants of interest (Eta (B.1.525), R.1, Zeta (P.2), Iota (B.1.526) and B.1.2/501Y or B.1.1.165). (**B**) ELISpot images and bar diagrams showing average frequencies of IFN-γ producing cell spots from immune cells from PBMCs (1 × 10^6^ cells per well) of COVID-19 infected with highly pathogenic SARS-CoV-2 variants of concern Beta (B.1.351) (*left panel*) and Omicron (B.1.1.529) (*right panel*). Cells were stimulated for 48 hours with 10mM of 16 immunodominant CD8^+^ T cell peptides derived from SARS-CoV-2 structural (Spike, Envelope, Membrane) and nonstructural (orf1ab, ORF6, ORF7b, ORF8a, ORF10) proteins. (**C**) ELISpot images and bar diagrams showing average frequencies of IFN-γ producing cell spots from immune cells from PBMCs (1 × 10^6^ cells per well) of COVID-19 infected with SARS-CoV-2 variants of concern Alpha (B.1.1.7) (*left panel*) and Omicron (B.1.1.529) (*right panel*). Cells were stimulated for 48 hours with 10mM of 6 immunodominant CD4^+^ T cell peptides derived from SARS-CoV-2 structural (Spike, Membrane, Nucleocapsid) and nonstructural (ORF1a, ORF6, ORF8a) proteins. The bar diagrams show the average/mean numbers (± SD) of IFN-γ-spot forming cells (SFCs) after CD8^+^ T cell peptide-stimulation PBMCs of Asymptomatic and Symptomatic COVID-19 patients. Dotted lines represent an arbitrary threshold set to evaluate the relative magnitude of the response. A strong response is defined for mean SFCs > 25 per 1 × 10^6^ stimulated PBMCs. Results were considered statistically significant at *P* < 0.05.

ASYMP COVID-19 patients showed significantly higher frequencies of SARS-CoV-2 epitope-specific IFN-γ-producing CD8^+^ T cells (mean SFCs > 25 per 1 × 10^6^ pulmonary immune cells), irrespective of infection with Beta (*P* < 0.5, **Fig. 1B**, *left panel*) or Omicron (*P* < 0., **Fig. 1B**, *right panel*) variants. In contrast, severely ill or hospitalized symptomatic COVID-19 patients showed significantly lower frequencies of SARS-CoV-2 epitope-specific IFN-γ-producing CD8^+^ T cells (*P* < 0.5, **Fig. 1B**, *left panel*) or Omicron (*P* < 0., **Fig. 1B**, *right panel*) variants. This observation was consistent regardless of whether CD8^+^ T cell’s targeted epitopes were from structural or non-structural SARS-CoV-2 protein antigens. suggesting that strong CD8^+^ T cell responses specific to selected “universal” SARS-CoV-2 epitopes were commonly associated with better COVID-19 outcomes. In contrast, low SARS-CoV-2-specific CD8^+^ T cell responses were more commonly associated with severe onset of disease.

Similarly, we found higher frequencies of functional IFN-γ-producing CD4^+^ T cells ASYMP COVID-19 patients (mean SFCs > 25 per 1 × 10^6^ pulmonary immune cells), irrespective of infection with Beta (*P* < 0.5, **Fig. 1C**, *left panel*) or Omicron (*P* < 0., **Fig. 1C**, *right panel*) variants. Whereas reduced frequencies of IFN-γ-producing CD4^+^ T cells were detected in SYMP COVID-19 patients, irrespective of infection with Beta (*P* < 0.5, **Fig. 1C**, *left panel*) or Omicron (*P* < 0., **Fig. 1C**, *right panel*) variants. This observation was consistent regardless of whether CD4^+^ T cell’s targeted epitopes were from structural or non-structural SARS-CoV-2 protein antigens. Our results suggest strong CD4^+^ T cell responses specific to selected “universal” SARS-CoV-2 epitopes were commonly associated with better COVID-19 outcomes. In contrast, low SARS-CoV-2-specific CD4^+^ T cell responses were more commonly associated with severe disease onset.

Altogether these results: (1) demonstrate an important role of SARS-CoV-2-specific CD4^+^ and CD8^+^ T cells directed against highly conserved structural and non-structural SARS-CoV-2 epitopes in protection from severe COVID-19 symptoms, and (2) highlight the potential importance of these highly conserved “asymptomatic” epitopes in mounting protected CD4^+^ and CD8^+^ T cell responses against multiple SARS-CoV-2 VOCs.

### 2. A pan-Coronavirus vaccine composed of a mixture of conserved “asymptomatic” CD4^+^ and CD8^+^ T cell epitopes provides robust protection against infection and disease caused by six SARS-CoV-2 variants of concern

We next used a prototype pan-Coronavirus vaccine composed of a mixture of 6 conserved “asymptomatic” CD4^+^ T cell epitopes and 16 conserved “asymptomatic” CD4^+^ and CD8^+^ T cell epitopes, previously identified that span the whole SARS-CoV-2 genome (33). We focused mainly on CD4^+^ and CD8^+^ T cell epitopes that show immunodominance selectively in SYMP COVID-19 patients infected with various SARS-CoV-2 VOCs.

A pool of peptides comprising 25_μ_g each of 16 CD8^+^ T cell peptides (ORF1ab_2210-2218_, ORF1ab_3013-3021_, ORF1ab_4283-4291_, ORF1ab_6749-6757_, ORF6_3-11_, ORF7b_26-34_, ORF8a_73-81_, ORF10_3-11_, ORF10_5-13_, S_958-966_, S_1000-1008_, S_1220-1228_, E_20-28_, E_26-34_, M_52-60_, and M_89-97_), 6 CD4^+^ T cell epitopes (ORF1a_1350-1365_, ORF6_12-26_, ORF8b_1-15_, S_1-13_, M_176-190_, and N_388-403_), and 7 B-cell peptides selected from the Spike protein, were mixed with cpG1826 adjuvant and administered subcutaneously on Day 0 and Day 14 to 7-8 week old triple transgenic HLA-A*02:01/HLA-DR hACE-2 mice (*n* = 30). The remaining group of the mock-immunized received vehicle alone (*n* = 30) (**Fig. 2A**). Fourteen days after the second immunization (i.e. day 28) mice were divided into 6 groups and intranasally infected with 1 × 10^5^ pfu of SARS-CoV-2 (USA-WA1/2020) (*n* = 10), 6 × 10^3^ pfu of SARS-CoV-2-Alpha (B.1.1.7) (*n* = 10), 6 × 10^3^ pfu of SARS-CoV-2-Beta (B.1.351) (*n* = 10), 5 × 10^2^ pfu of SARS-CoV-2-Gamma (P.1) (*n* = 10), 8 × 10^3^ pfu of SARS-CoV-2-Delta (B.1.617.2) (*n* = 10), and 6.9 × 10^4^ pfu of SARS-CoV-2-Omicron (B.1.1.529) (*n* = 10) (**Fig. 2A**).

**Figure 2.**
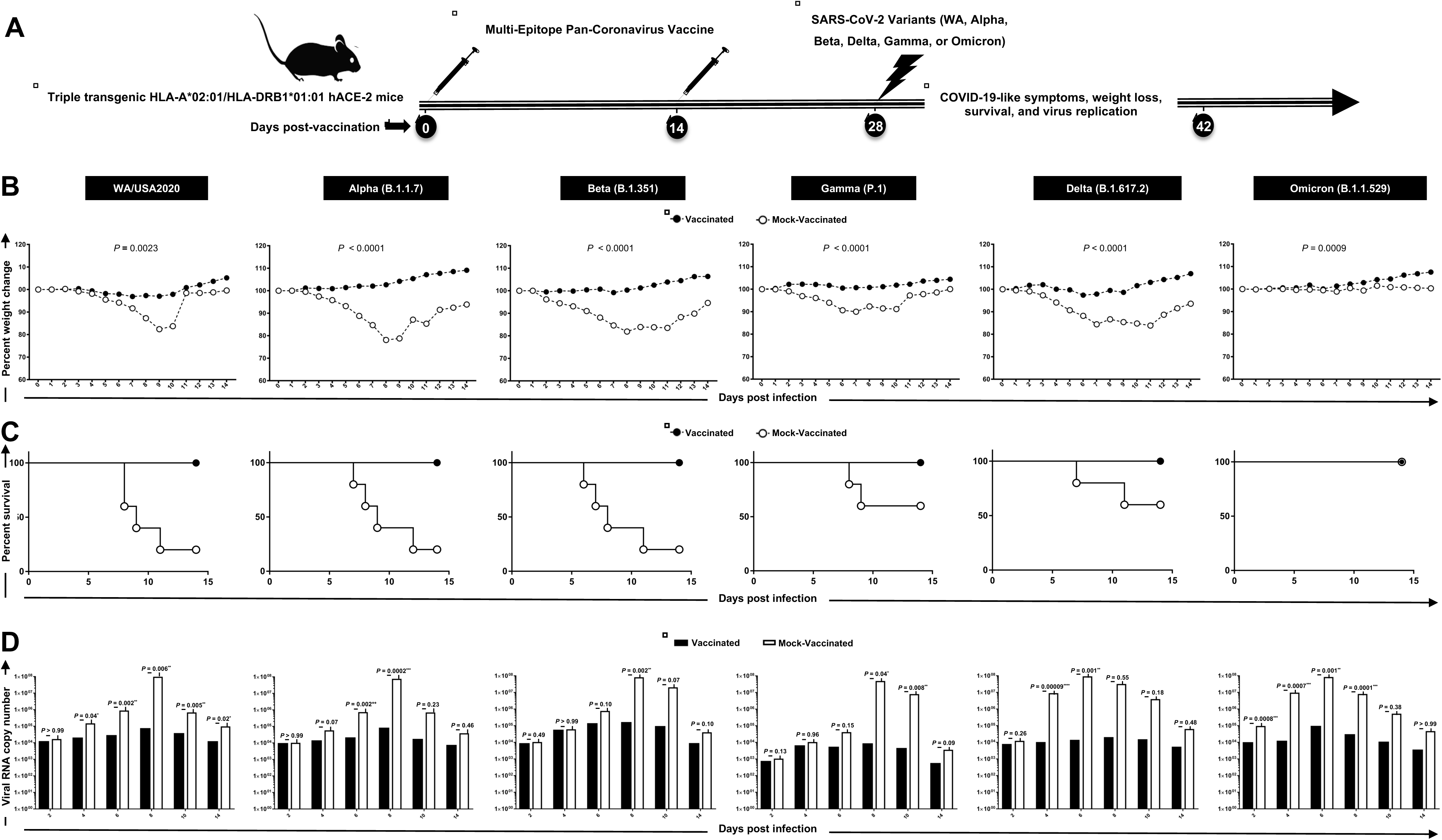
Protection induced against six SARS-CoV-2 variants of concern in triple transgenic HLA-A*02:01/HLA-DRB1*01:01-hACE-2 mice following immunization with a pan-Coronavirus vaccine incorporating conserved human B, CD4^+^, and CD8^+^ T cell “asymptomatic” epitopes: (**A**) Experimental scheme of vaccination and challenge triple transgenic HLA-A*02:01/HLA-DRB1*01:01-hACE-2 mice. Triple transgenic HLA-A*02:01/HLA-DRB1*01:01-hACE-2 mice (7-8-week-old, *n* = 60) were immunized subcutaneously on Days 0 and 14 with a multi-epitope pan-Coronavirus vaccine consisting of a pool of conserved B, CD4^+^ T cell and CD8^+^ T cell human epitope peptides. The pool of peptides comprised 25_μ_g of each of the 16 CD8^+^ T cell peptides, 6 CD4^+^ T cell peptides, and 7 B-cell peptides. The final composition of peptides was mixed with 25_μ_g of CpG and 25_μ_g of Alum. Mock-vaccinated mice were used as controls (*Mock*). Fourteen days following the second immunization, mice were intranasally challenged with each of the six different SARS-CoV-2 variants of concern (WA/USA2020, Alpha (B.1.1.7), Beta (B.1.351), Gamma (P.1), Delta (B.1.617.2), and Omicron (B.1.1.529)). Vaccinated and mock-vaccinated mice were followed 14 days post-challenge for COVID-like symptoms, weight loss, survival, and virus replication. (**B**) Percent weight change recorded daily for 14 days p.i. in vaccinated and mock-vaccinated mice following the challenge with each of the six different SARS-CoV-2 variants. (**C**) Kaplan-Meir survival plots for vaccinated and mock-vaccinated mice following the challenge with each of the six different SARS-CoV-2 variants. (**D**) Virus replication in vaccinated and mock-vaccinated mice following the challenge with each of the six different SARS-CoV-2 variants detected in throat swabs on Days 2, 4, 6, 8, 10, and 14, The indicated *P* values are calculated using the unpaired *t*-test, comparing results obtained in vaccinated VERSUS mock-vaccinated mice.

Mice that received the pan-Coronavirus vaccine showed significant protection from weight loss (**Fig. 2B**) and death (**Fig. 2C**) following infection with each of the six SARS-CoV-2 variants of concern: WA/USA2020, Alpha (B.1.1.7), Beta (B.1.351), Gamma (P.1), Delta (B.1.617.2), and Omicron (B.1.1.529). All mice immunized with the conserved pan-Coronavirus vaccine survived infection with SARS-CoV-2 variants of concern. In contrast to mock-immunized mice where 60% mortality was detected among WA/USA2020 infected mice, 80% mortality among Alpha (B.1.1.7) and Beta (B.1.351) infected mice, 40% mortality among Gamma (P.1) and Delta (B.1.617.2) variants infected mice (**Fig. 2C**). Mortality was not observed for mock-immunized mice infected with the SARS-CoV-2 Omicron (B.1.1.529) variant (**Fig. 2C**).

Throat swabs were collected from the vaccinated and mock-vaccinated groups of mice on days 2, 4, 6, 8, 10, and 14 post-infection (p.i.) and were processed to detect the viral RNA copy number by qRT-PCR (**Fig. 2D**). Compared to the viral RNA copy number detected from the mock-vaccinated group of mice, we detected a statistically significant decrease in the viral RNA copy number among vaccinated groups of mice on day 4 p.i. for SARS-CoV-2 WA/USA2020 (*P* = 0.04), Delta (B.1.617.2) (*P* = 0.00009), and Omicron (B.1.1.529) (*P* = 0.007); on day 6 p.i. for SARS-CoV-2 WA/USA2020 (*P* = 0.002), Alpha (B.1.1.7) (*P* = 0.002), Delta (B.1.617.2) (*P* = 0.001), and Omicron (B.1.1.529) (P = 0.001); on day 8 p.i. for SARS-CoV-2 WA/USA2020 (*P* = 0.006), Alpha (B.1.1.7) (*P* = 0.0002), Beta (B.1.351) (*P* = 0.002), Gamma (P.1) (*P* = 0.04), and Omicron (B.1.1.529) (*P* = 0.0001); on day 10 p.i. for SARS-CoV-2 WA/USA2020 (*P* = 0.005), Gamma (P.1) (*P* = 0.008); and on day 14 p.i. for SARS-CoV-2 WA/USA2020 (*P* = 0.02) (**Fig. 2D**). This result suggests that the pan-Coronavirus vaccine showed significant protection from virus replication for most of SARS-CoV-2 variants and confirms a plausible anti-viral effect following immunization with asymptomatic B, CD4^+^ and CD8^+^ T cell epitopes carefully selected as being highly conserved from multiple SARS-CoV-2 variants.

### 3. Immunization with the Pan-Coronavirus vaccine bearing conserved epitopes reduced COVID-19-related lung pathology and virus replication associated with increased infiltration of CD8^+^ and CD4^+^ T cells in the lungs

Hematoxylin and eosin staining of lung sections at day 14 p.i. showed a significant reduction in COVID-19-related lung pathology in the mice immunized with conserved Pan-Coronavirus vaccine compared to mock-vaccinated mice (**Fig. 3A**). This reduction in lung pathology was observed for all six SARS-CoV-2 variants: USA-WA1/2020, Alpha (B.1.1.7), Beta (B.1.351), Gamma (P.1), Delta (B.1.617.2), and Omicron (B.1.1.529) showed severe lung pathogenicity (**Fig. 3A**). We further performed SARS-CoV-2 Nucleocapsid Antibody-based Immunohistochemistry (IHC) staining on lung tissues obtained from vaccinated and mock-vaccinated groups of mice infected with SARS-CoV-2 variants. We detected significantly lower antibody staining in the lung tissues of the vaccinated compared mock-vaccinated group of mice following infection with each of the six SARS-CoV-2 variants of concern. This indicated higher expression of the target viral proteins in the lungs of the mock-vaccinated compared to the vaccinated group of mice (**Fig. 3B**). Furthermore, IHC staining was performed to compare the infiltration CD8^+^ and CD4^+^ T cells into lung tissues of vaccinated and mock-vaccinated mice infected with various SARS-CoV-2 variants. Forten days following infection with each of the six variants, we observed a significant increase in the infiltration of both CD8^+^ T cells (**Fig. 3C**) and CD4^+^ T cells (**Fig. 3D**) in the lungs of vaccinated mice compared to mock-vaccinated mice.

**Figure 3.**
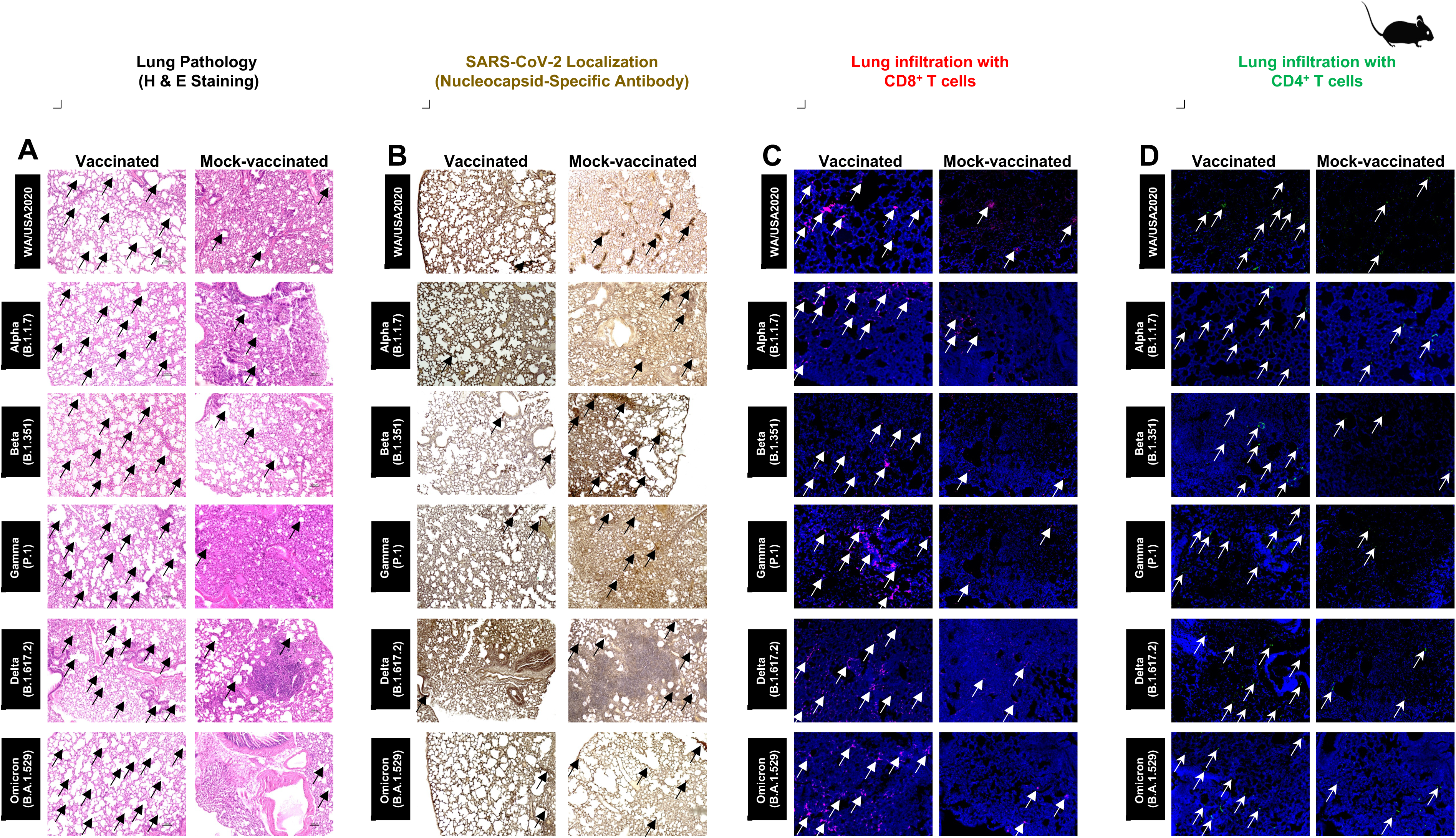
Histopathology and immunohistochemistry of the lungs from in triple transgenic HLA-A*02:01/HLA-DRB1*01:01-hACE-2 mice vaccinated and mock-vaccinated mice. (**A**) Representative images of hematoxylin and Eosin (H & E) staining of the lungs harvested on day 14 p.i. from vaccinated (*left panels*) and mock-vaccinated (*right panels*) mice. (**B**) Representative immunohistochemistry (IHC) sections of the lungs were harvested on Day 14 p.i. from vaccinated (*left panels*) and mock-vaccinated (*right panels*) mice and stained with SARS-CoV-2 Nucleocapsid antibody. Black arrows point to the antibody staining. Fluorescence microscopy images showing infiltration of CD8^+^ T cells (**C**) and of CD4^+^ T cells (**D**) in the lungs from vaccinated (*left panels*) and mock-vaccinated (*right panels*) mice. Lung sections were co-stained using DAPI (*blue*) and mAb specific to CD8^+^ T cells (*Pink*) (magnification, 20x). The white arrows point to CD8^+^ and CD4^+^ T cells infiltrating the infected lungs.

Altogether these results indicate that immunization with the Pan-Coronavirus vaccine bearing conserved epitopes induced cross-protective CD8^+^ and CD4^+^ T cells that infiltrated the lungs cleared the virus, and reduced COVID-19-related lung pathology following infection with various multiple SARS-CoV-2 variants.

### 4. Increased frequencies of lung-resident functional CD8^+^ and CD4^+^ T_EM_ and T_RM_ cells induced by the Pan-Coronavirus vaccine are associated with protection against multiple SARS-CoV-2 variants

To determine whether increased frequencies of lung-resident functional CD8^+^ and CD4^+^ T cells induced by the pan-Coronavirus vaccine are associated with protection against multiple SARS-CoV-2 variants, we used flow cytometry and compared the frequencies of IFN-γ CD8^+^ T cells and CD69 CD8^+^ T cells (**Fig. 4A**), IFN-γ CD4^+^ T cells and CD69 CD4^+^ T cells (**Fig. 4B**) in cell suspensions from the lungs of vaccinated versus mock-vaccinated groups of mice.

**Figure 4.**
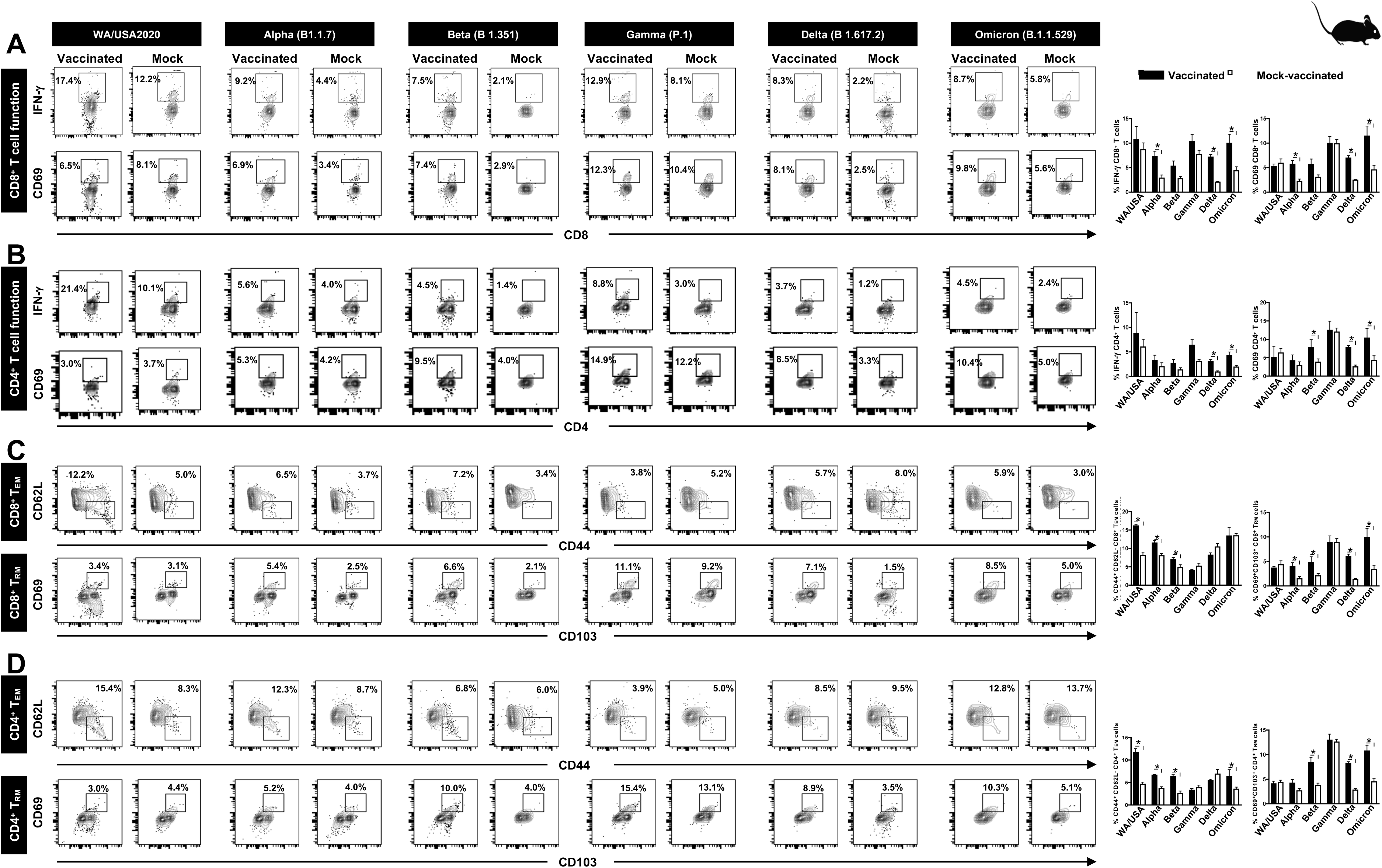
The effect of Pan-Coronavirus immunization on CD8^+^ and CD4^+^ T cell function and memory response: FACS plots and bar graphs showing the (**A**) expression of CD8^+^ T cell function markers, (**B**) CD4^+^ T cell function associated markers, (**C**) CD8^+^ T effector memory response (CD44^+^CD62L^−^), and CD8^+^ T resident memory (CD103^+^CD69^+^) response, and (**D**) CD4^+^ T effector memory response (CD44^+^CD62L^−^), and CD4^+^ resident memory (CD103^+^CD69^+^) response in the lung of vaccinated and mock-vaccinated groups of mice infected with multiple SARS-CoV-2 variants. Bars represent means ± SEM. Data were analyzed by student’s *t*-test. Results were considered statistically significant at *P* < 0.05.

Relatively higher frequencies of IFN-γ CD8^+^ T cells were detected in the lungs of protected mice that received the pan-Coronavirus vaccine compared to non-protected mock-vaccinated mice following infections with various SARS-CoV-2 variants: USA-WA1/2020 (Vaccinated = 17.4% vs. Mock = 12.2%, *P* = 0.5178), Alpha (B.1.1.7) (Vaccinated = 9.2% vs. Mock = 4.4%, *P* = 0.0076), Beta (B.1.351) (Vaccinated = 7.5% vs Mock = 2.1%, *P* = 0.05), Gamma (P.1) (Vaccinated = 12.9% vs. Mock = 8.1%, *P* = 0.14), Delta (B.1.617.2) (Vaccinated = 8.3% vs. Mock = 2.23%, *P* < 0.0001), and Omicron (B.1.1.529) (Vaccinated = 8.7% vs. Mock = 5.8%, *P* = 0.02) (**Fig. 4A**, top row). Similarly, increased frequencies for CD8^+^CD69^+^ T cells were detected in the lungs of protected mice that received the pan-Coronavirus vaccine compared to non-protected mock-vaccinated mice following infections with various SARS-CoV-2 variants: Alpha (B.1.1.7) (Vaccinated = 6.9% vs Mock = 3.4%, *P* = 0.0033), Beta (B.1.351) (Vaccinated = 7.4% vs Mock = 2.9%, *P* = 0.05), Gamma (P.1) (Vaccinated = 12.3% vs Mock = 10.4%, *P* = 0.95), Delta (B.1.617.2) (Vaccinated = 8.1% vs Mock = 2.5%, *P* < 0.0001), and Omicron (B.1.1.529) (Vaccinated = 9.8% vs Mock = 5.6%, *P* = 0.01) (**Fig. 4A**, bottom row).

Moreover, higher frequencies of IFN-γ CD4^+^ T cells were detected in the lungs of protected mice that received the pan-Coronavirus vaccine compared to non-protected mock-vaccinated mice following infections with various SARS-CoV-2 variants: USA-WA1/2020 (Vaccinated = 21.4% vs Mock = 10.1%, *P* = 0.5696), Alpha (B.1.1.7) (Vaccinated = 5.6% vs Mock = 4%, *P* = 0.35), Beta (B.1.351) (Vaccinated = 4.5% vs Mock = 1.4%%, *P* = 0.12), Gamma (P.1) (Vaccinated = 8.8% vs Mock = 3%, *P* = 0.02), Delta (B.1.617.2) (Vaccinated = 3.7% vs Mock = 1.2%, *P* = 0.0002), and Omicron (B.1.1.529) (Vaccinated = 4.5% vs Mock = 2.4%, *P* = 0.01) (**Fig. 4B**, top row). Similarly, increased frequencies for CD4^+^CD69^+^ T cells were detected in the lungs of protected mice that received the pan-Coronavirus vaccine compared to non-protected mock-vaccinated mice following infections with various SARS-CoV-2 variants: Alpha (B.1.1.7) (Vaccinated = 5.3% vs Mock = 4.2%, *P* = 0.1748), Beta (B.1.351) (Vaccinated = 9.5% vs Mock = 4%, *P* = 0.009), Gamma (P.1) (Vaccinated = 14.9% vs Mock = 12.2%, *P* = 0.7155), Delta (B.1.617.2) (Vaccinated = 8.5% vs Mock = 3.3%, *P* < 0.0001), and Omicron (B.1.1.529) (Vaccinated = 10.4% vs Mock = 5%, *P* = 0.003) (**Fig. 4B**, bottom row).

FACS-based immunophenotyping, confirmed higher frequencies of the memory CD8^+^ T_EM_ (CD44^+^CD62L^−^) cell subset in immunized mice with a pool of pan-Coronavirus peptides and subjected to infection against USA-WA1/2020 (Vaccinated = 12.2% vs Mock = 5%, P < 0.0001), Alpha (B.1.1.7) (Vaccinated = 6.5% vs Mock = 3.7%, P = 0.0017), Beta (B.1.351) (Vaccinated = 7.2% vs Mock = 3.4%, P = 0.0253), and Omicron (B.1.1.529) (Vaccinated = 5.9% vs Mock = 3%, P = 0.9765) (**Fig. 4C**). Similarly, when the frequencies for the memory CD8^+^ T_RM_ (CD69^+^CD103^+^) cell subset was evaluated, we found higher CD8^+^ T_RM_ cell subset frequencies for immunized mice infected with USA-WA1/2020 (Vaccinated = 3.4% vs Mock = 3.1%, P = 0.4004), Alpha (B.1.1.7) (Vaccinated = 5.4% vs Mock = 2.5%, P = 0.0160), Beta (B.1.351) (Vaccinated = 6.6% vs Mock = 2.1%, P = 0.0420), Gamma (P.1) (Vaccinated = 11.1% vs Mock = 9.2%, P = 0.9961), Delta (B.1.617.2) (Vaccinated = 7.1% vs Mock = 1.5%, P < 0.0001), and Omicron (B.1.1.529) (Vaccinated = 8.5% vs Mock = 5%, P = 0.0139) (**Fig. 4C**).

Moreover, in context to memory CD4^+^ T_EM_ (CD44^+^CD62L^−^) cell subset, relatively higher frequencies were observed for immunized mice subjected to infection with SARS-CoV-2 variants USA-WA1/2020 (Vaccinated = 15.4% vs Mock = 8.3%, P = 0.0001), Alpha (B.1.1.7) (Vaccinated = 12.3% vs Mock = 8.7%, P < 0.0001), and Beta (B.1.351) (Vaccinated = 6.8% vs Mock = 6%, P < 0.0004) (**Fig. 4D**). Higher frequencies of the CD4^+^ T_RM_ (CD69^+^CD103^+^) cell subset were found in immunized mice infected with SARS-CoV-2 variants Alpha (B.1.1.7) (Vaccinated = 5.2% vs Mock = 4%, P = 0.0828), Beta (B.1.351) (Vaccinated = 10% vs Mock = 4%, P = 0.005), Gamma (P.1) (Vaccinated = 15.4% vs Mock = 13.1%, P = 0.7860), Delta (B.1.617.2) (Vaccinated = 8.9% vs Mock = 3.5%, P < 0.0001), and Omicron (B.1.1.529) (Vaccinated = 10.3% vs Mock = 5.1%, P = 0.0021) (**Fig. 4D**).

Altogether, our findings confirmed that immunization with the pan-Coronavirus vaccine bearing conserved epitopes induced high frequencies of functional CD8^+^ and CD4^+^ T_EM_ and T_RM_ cells that infiltrate the lungs associated with a significant decrease in virus replication and a reduction in COVID-19-related lung pathology following infection with various multiple SARS-CoV-2 variants.

### 5. Increased SARS-CoV-2 epitopes-specific IFN-γ-producing CD8^+^ T cells in the lungs of vaccinated mice in comparison to mock-vaccinated mice

To determine whether the functional lung-resident CD8^+^ T cells are specific to SARS-CoV-2, we stimulated lung-cell suspension from vaccinated and mock-vaccinated mice with each of the 14 “universal” human CD8^+^ T cell epitopes (ORF1ab_2210-2218_, ORF1ab_3013-3021_, ORF1ab_4283-4291_, ORF1ab_6749-6757_, ORF6_3-11_, ORF7b_26-34_, ORF10_3-11_, ORF10_5-13_, S_958-966_, S_1000-1008_, S_1220-1228_, E_20-28_, M_52-60_, and M_89-97_) and quantified the number of IFN-γ-producing CD8^+^ T cells using ELISpot, as detailed in the *Materials & Methods* section (**Fig. 5**). To determine where cross-reactive IFN-γ-producing CD8^+^ T cell responses will be detected regardless of SARS-CoC-2 variant, the number IFN-γ-producing CD8^+^ T cells were determined in the lung tissues of vaccinated and mock-vaccinated mice after challenge with each of six different SARS-CoV-2 variants of concern.

**Figure 5.**
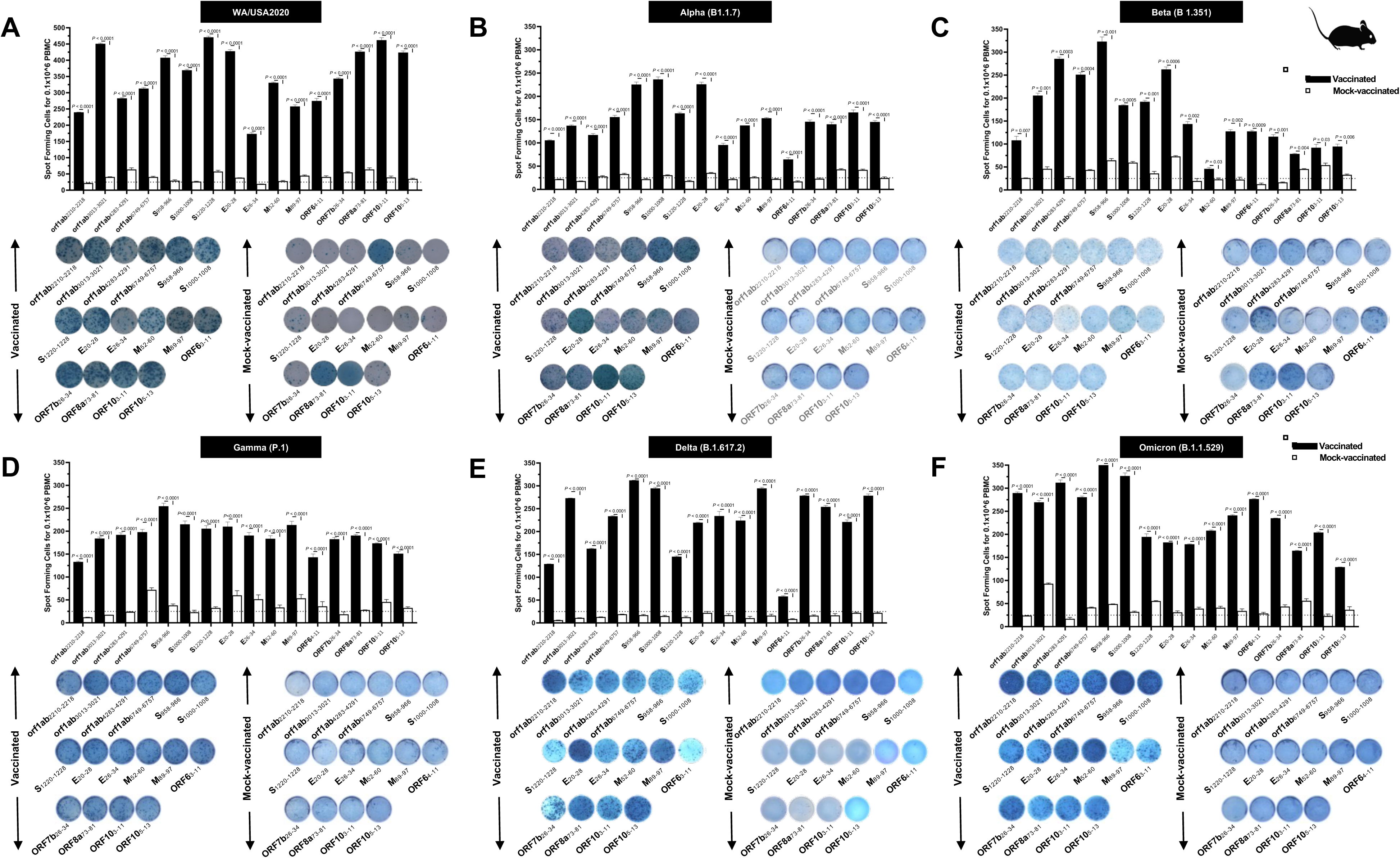
Immunogenicity of conserved SARS-CoV-2 CD8^+^ T cell epitopes in triple transgenic HLA-A*02:01/HLA-DRB1*01:01-hACE-2 mice: ELISpot images and bar diagrams showing average frequencies of IFN-γ producing cell spots from mononuclear cells from lung tissue (1 × 10^6^ cells per well) of vaccinated and mock-vaccinated mice challenged with (**A**) WA/USA2020, (**B**) Alpha (B.1.1.7), (**C**) Beta (B.1.351), (**D**) Gamma (P.1), (**E**) Delta (B.1.617.2), and (**F**) Omicron (B.1.1.529). The cells were stimulated for 48 hours with 10mM of 16 immunodominant CD8^+^ T cell peptides. The bar diagrams show the average/mean numbers (± SD) of IFN-γ-spot forming cells (SFCs) after CD8^+^ T cell peptide stimulation in lung tissues of vaccinated and mock-vaccinated mice. Dotted lines represent an arbitrary threshold set to evaluate the relative magnitude of the response. A strong response is defined for mean SFCs > 25 per 1 × 10^6^ stimulated PBMCs. Results were considered statistically significant at *P* < 0.05.

Overall, a significant increase in the number of IFN-γ-producing CD8^+^ T cells was detected in the lungs of protected mice that received the pan-Coronavirus vaccine compared to non-protected mock-vaccinated mice (mean SFCs > 25 per 0.5 × 10^6^ pulmonary immune cells), irrespective of the SARS-CoV-2 variants of concern: WA/USA2020 (**Fig. 5A**), Alpha (B.1.1.7) (**Fig. 5B**), Beta (B.1.351) (**Fig. 5C**), Gamma (P.1) (**Fig. 5D**), Delta (B.1.617.2) (**Fig. 5E**), or Omicron (B.1.1.529) (**Fig. 5F**). All the comparisons among vaccinated and mock-vaccinated groups of mice, irrespective of SARS-CoV-2 variants of concern were found to be statistically significant regardless of whether CD8^+^ T cells targeted epitopes were from structural (Spike, Envelope, Membrane), or non-structural (ORF1ab, ORF6, ORF7b, ORF10) SARS-CoV-2 protein antigens (*P* < 0.5).

Taken together, these results: (1) Confirm that immunization with the pan-Coronavirus vaccine bearing conserved epitopes induced high frequencies of functional CD8^+^ T cells that infiltrate the lungs associated with cross-protection against multiple SARS-CoV-2 variants; (2) Demonstrate that increased SARS-CoV-2 epitopes-specific IFN-γ-producing CD8^+^ T cells in the lungs of vaccinated triple transgenic HLA-A*02:01/HLA-DRB1*01:01-hACE-2 mice are associated with protection from multiple variants of concern. In contrast, low frequencies of lung-resident SARS-CoV-2-specific IFN-γ-producing CD8^+^ T cells were associated with severe disease onset in mock-vaccinated triple transgenic HLA-A*02:01/HLA-DRB1*01:01-hACE-2 mice. In this report, we suggest an important role for functional lung-resident SARS-CoV-2-specific CD8^+^ T cells specific to highly conserved “universal” epitopes from structural and non-structural antigens in cross-protection against SARS-CoV-2 VOCs.

### 6. Increased SARS-CoV-2 epitopes-specific IFN-γ-producing CD4^+^ T cells in the lungs of vaccinated mice in comparison to mock-vaccinated mice

We stimulated lung-cell suspension from vaccinated and mock-vaccinated groups of mice with each of the 6 “universal” human CD4^+^ T cell epitopes (ORF1a_1350-1365_, ORF6_12-26_, ORF8b_1-15_, S_1-13_, M_176-190_, and N_388-403_) and quantified the number of IFN-γ-producing CD4^+^ T cells using ELISpot, to determine whether the functional lung-resident CD4^+^ T cells are specific to SARS-CoV-2 (**Fig. 6**).

**Figure 6.**
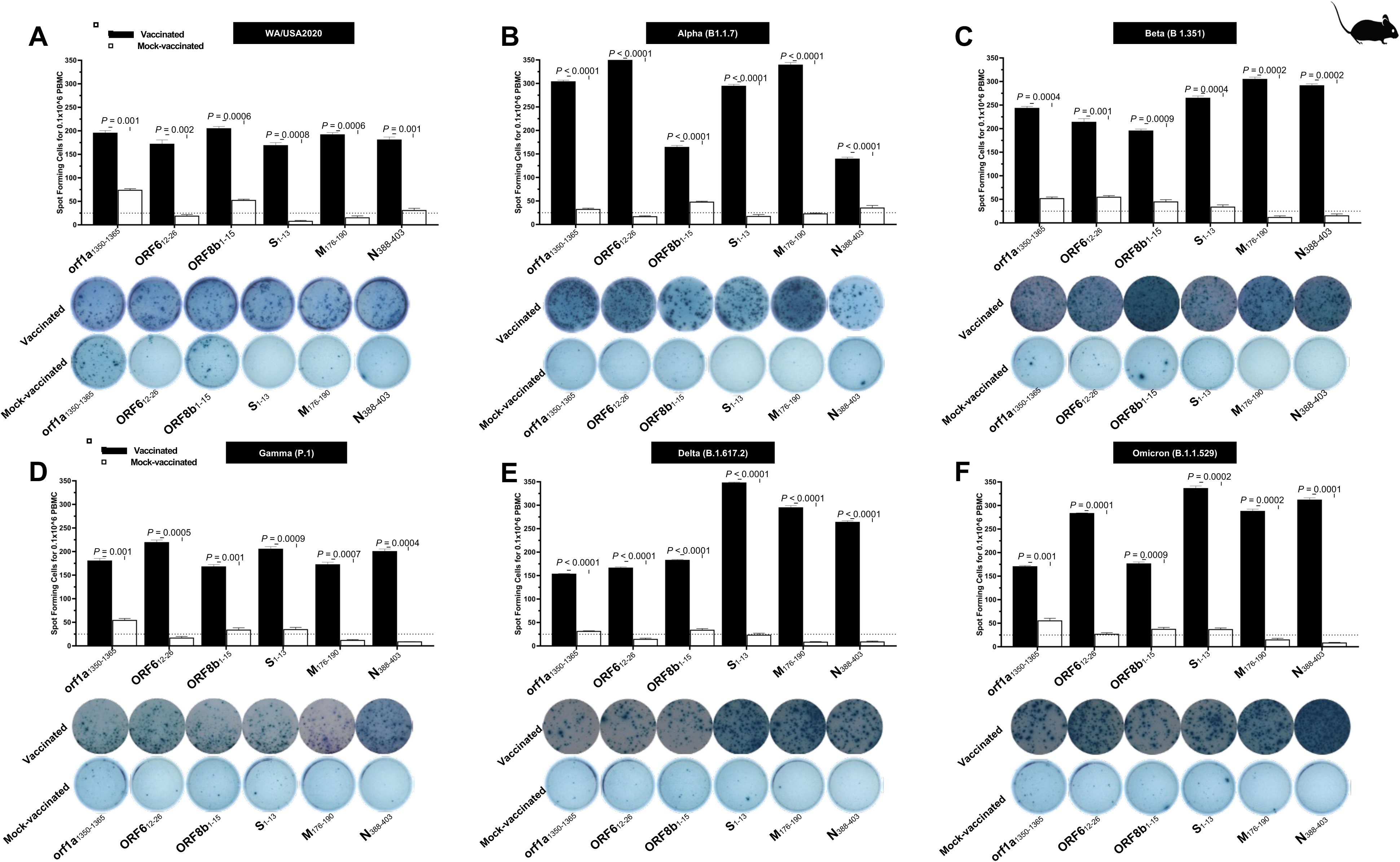
The magnitude of the IFN-γ CD4^+^ T cell responses for 6 conserved SARS-CoV-2 CD4^+^ T cell epitopes in triple transgenic HLA-A*02:01/HLA-DRB1*01:01-hACE-2 mice: ELISpot images and bar diagrams showing average frequencies of IFN-γ producing cell spots from mononuclear cells from lung tissue (1 × 10^6^ cells per well) of vaccinated and mock-vaccinated mice challenged with (**A**) WA/USA2020, (**B**) Alpha (B.1.1.7), (**C**) Beta (B.1.351), (**D**) Gamma (P.1), (**E**) Delta (B.1.617.2), and (**F**) Omicron (B.1.1.529). Cells were stimulated for 48 hours with 10mM of 6 immunodominant CD4^+^ T cell peptides derived from SARS-CoV-2 structural (Spike, Envelope, Membrane) and nonstructural (orf1ab, ORF6, ORF7b, ORF8a, ORF10) proteins. The bar diagrams show the average/mean numbers (± SD) of IFN-γ-spot forming cells (SFCs) after CD8^+^ T cell peptide stimulation in lung tissues of vaccinated and mock-vaccinated mice. The dotted lines represent an arbitrary threshold set to evaluate the relative magnitude of the response. A strong response is defined for mean SFCs > 25 per 1 × 10^6^ stimulated PBMCs. Results were considered statistically significant at *P* ≤ 0.05.

Overall, we detected a significant increase in the number of IFN-γ-producing CD4^+^ T cells in the lungs of protected mice that received the pan-Coronavirus vaccine compared to non-protected mock-vaccinated mice (mean SFCs > 25 per 0.5 × 10^6^ pulmonary immune cells), irrespective of the SARS-CoV-2 VOCs: WA/USA2020 (**Fig. 6A**), Alpha (B.1.1.7) (**Fig. 6B**), Beta (B.1.351) (**Fig. 6C**), Gamma (P.1) (**Fig. 6D**), Delta (B.1.617.2) (**Fig. 6E**), or Omicron (B.1.1.529) (**Fig. 6F**). All the comparisons among vaccinated and mock-vaccinated groups of mice, irrespective of SARS-CoV-2 VOCs were statistically significant regardless of whether CD4^+^ T cells targeted epitopes were from structural or non-structural SARS-CoV-2 protein antigens (*P* < 0.5).

Taken together, our findings demonstrate that increased SARS-CoV-2 epitopes-specific IFN-γ-producing CD4^+^ T cells in the lungs of vaccinated triple transgenic HLA-A*02:01/HLA-DRB1*01:01-hACE-2 mice are associated with protection from multiple variants of concern. In contrast, low frequencies of lung-resident SARS-CoV-2-specific IFN-γ-producing CD4^+^ T cells were associated with severe disease onset in mock-vaccinated triple transgenic HLA-A*02:01/HLA-DRB1*01:01-hACE-2 mice. The findings suggest an important role of functional lung-resident SARS-CoV-2-specific CD4^+^ T cells specific to highly conserved “universal” epitopes from structural and non-structural antigens in cross-protection against SARS-CoV-2 VOCs.

### 7. Universal B cell epitopes from SARS-CoV-2 Spike protein showed a high degree of immunogenicity across SARS-CoV-2 variants based on antibody response in COVID-19 patients and triple transgenic HLA-A*02:01/HLA-DRB1*01:01-hACE-2

We next determined whether the antibody responses were associated with protection since the prototype pan-Coronavirus vaccine used herein also contains nine conserved B cell epitopes selected from the Spike glycoprotein of SARS-CoV-2. The nine B-cell epitopes were screened for their conservancy against variants namely h-CoV-2/Wuhan (MN908947.3), h-CoV-2/WA/USA2020 (OQ294668.1), h-CoV-2/Alpha(B1.1.7) (OL689430.1), h-CoV-2/Beta(B 1.351) (MZ314998), h-CoV-2/Gamma(P.1) (MZ427312.1), h-CoV-2/Delta(B.1.617.2) (OK091006.1), and h-CoV-2/Omicron(B.1.1.529) (OM570283.1). We observed 100% conservancy in three of our earlier predicted B cell epitopes namely S_287-317_, S_524-558_, and S_565-598_ (**Fig. S3**).

The antibody titer specific to each of the nine “universal” B-cell epitopes was determined by ELISA in COVID-19 patients infected with multiple SARS-CoV-2 variants of concern (**Fig. S4**, *left panel*) and in vaccinated and mock-vaccinated triple transgenic HLA-A*02:01/HLA-DRB1*01:01-hACE-2 mice challenged with same SARS-CoV-2 VOCs (**Fig. S4**, *right panel*). The peptide binding IgG level was significantly higher for all nine “universal” B cell epitopes in COVID-19 patients (**Fig. S4**, *left panel*) as well as in vaccinated triple transgenic mice (**Fig. S4**, *right panel*), irrespective of SARS-CoV-2 variant. Reduced peptide binding IgG level was observed for severely ill COVID-19 patients (**Fig. S4**, *left panel*) and in mock-vaccinated triple transgenic HLA-A*02:01/HLA-DRB1*01:01-hACE-2 mice (**Fig. S4**, *right panel*).

Altogether, these results indicate that immunization with the pan-Coronavirus vaccine bearing conserved “universal” B and T cell epitope induced cross-protective antibodies, CD8^+^ and CD4^+^ T cells that infiltrated the lungs, cleared the virus, and reduced COVID-19-related lung pathology following infection with various multiple SARS-CoV-2 VOCs.

## DISCUSSION

Current Spike-based COVID-19 vaccines have contributed to a significantly decreased rate of SARS-CoV-2 infections. However, the long-term outlook of COVID-19 remains a serious cause of high death worldwide; with the mortality rate still surpassing even the worst mortality rates recorded for the influenza viruses. The continuous emergence of SARS-CoV-2 variants and sub-variants of concern, including the recent heavily mutated and highly transmissible Omicron sub-variants, has led to vaccine breakthroughs that contributed to prolonging the COVID-19 pandemic. The decrease over time in neutralizing antibody titers induced by current Spike-based vaccines, along with these vaccine breakthrough infections due to mutations on the Spike protein in recent variants and sub-variants, point to the urgent need to develop a next-generation B- and T-cell-based pan-Coronavirus vaccine-coronavirus vaccine, that would be based not only on Spike protein but also on less-mutated non-Spike structural and non-structural antigens and epitopes. Such a universal CoV vaccine could induce strong and durable protective immunity against infections and diseases caused by multiple emerging SARS-CoV-2 variants and sub-variants.

Much of the data on the efficacy of the current modified messenger RNA (mRNA) vaccines has shown that these vaccines elicited lower levels of neutralizing antibodies against newer SARS-CoV-2 variants than against the older variants. In the present report, we have identified “universal” CD8^+^ & CD4^+^ T cell and B cell epitopes conserved among all known SARS-CoV-2 variants, previous SARS and MERS coronavirus strains, and strains specific to different species which were reported to be hosts for SARS/MERS (bat, civet cat, pangolin, camel). The screening of these “universal” epitopes is limited to the spike alone and all the remaining structural and non-structural proteins of SARS-CoV-2. We used a combination of these highly conserved CD8^+^ & CD4^+^ T cell and B cell epitopes to design our first multi-epitope pan-Coronavirus vaccine.

We demonstrated that immunization of triple transgenic h-ACE-2-HLA-A2/DR mice with a pool of “universal” CD8^+^ T cell, CD4^+^ T cell, and B cell peptides produced protection against 6 variants including Washington, Alpha (B.1.1.7), Beta (B.1.351), Gamma (P.1), Delta (B.1.617.2), and Omicron (B.1.1.529), variants of SARS-CoV-2. The Pan-Coronavirus vaccine was found to be safe, as no local or systemic side effects were observed in the vaccinated mice. Moreover, we found that protection correlated with high frequencies of IFN-γ CD4^+^ T cells, CD69 CD4^+^ T cells, IFN-γ CD8^+^ T cells, and CD69 CD8^+^ T cells infuriating the lungs. We also found higher frequencies for the CD8^+^ T_EM_ (CD44^+^CD62L^−^) cell population in the lungs of protected mice. High levels of peptide-specific IgG were also detected in protected animals suggesting the contribution of Spike-specific antibodies in protection. A stark difference in the level of neutralizing viral titer was also observed between the vaccinated and mock-vaccinated groups of mice for all the studied variants. We observed no mortality in the vaccinated mice, irrespective of the SARS-CoV-2 variant. In contrast, high mortality was observed in the mock-vaccinated mice when challenged with 6 SARS-CoV-2 variants. Overall, the reported universal Coronavirus vaccine was safe, immunogenic, and provided cross-protection against multiple SARS-CoV-2 variants of Concern.

A typical SARS-CoV-2 virus accumulates 1-2 single-nucleotide mutations in its genome per month, which is ½ the rate of influenza and ¼ of the rate of HIV. Part of the reason that SARS-CoV-2 appears to be mutating more slowly is that, unlike most RNA viruses, coronaviruses have a novel exoribonuclease (ExoN) encoded in their genomes, which researchers suspect is correcting many of the errors that occur during replication. Genetic inactivation of this exonuclease in SARS-CoV and Murine coronavirus (MHV) increased mutation rates by 15 to 20-fold. The molecular basis of this CoV proofreading complex is being investigated as a possible therapeutic target for SARS-CoV-2. Importantly, nucleotide deletions, unlike substitutions, cannot be corrected by this proofreading mechanism, which is a factor that may accelerate adaptive evolution to some degree. Depending on the specific mutation, and where in the genome the nucleotide substitution, addition, or deletion occurs, mutations may be neutral, beneficial, or harmful to an organism. SARS-CoV-2’s spike (S) protein is 1273 amino acids long and is the main target of current COVID-19 vaccines, as well as those in development. It is the portion of the virus that recognizes and binds to host cellular receptors and mediates viral entry. SARS-CoV-2 is unable to infect host cells without it. Because of this, mutations in the S gene, particularly those that affect portions of the protein that are critical for pathogenesis and normal function (such as the receptor binding domain (RBD) or furin cleavage site) or those that cause conformational changes to the S protein, are of the greatest interest. If these changes are not recognized by “first-wave” antibodies, these mutations may provide an avenue for the virus to escape from immunity to the original SARS-CoV-2 strain.

The first reported SARS-CoV-2 mutation, D614G, which has now become common to nearly all sequenced SARS-CoV-2 genomes worldwide, followed by an analysis of additional key S protein mutations of several identified SARS-CoV-2 variants: B.1.1.7, commonly dubbed the U.K. variant; B.1.351, also known as 501Y.V2 or the South African variant, P.1., also known as 501Y.V3 or the Brazilian variant; B.1.427 and B.1.429, also recognized as CAL.20C or the California variant; B.1.526, or the New York variant and multiple lineages of variants that contain mutations at amino acid position 677.

Initial reports that a mutation had been identified in the SARS-CoV-2 genome began circulating in March 2020, and by the end of June, D614G, which constitutes the replacement of aspartate (D) with glycine (G) at the 614th amino acid of S protein, was found in nearly all SARS-CoV-2 samples worldwide. D614G has been found to enhance viral replication in human lung epithelial cells and primary human airway tissues by increasing the infectivity and stability of virions. Additional research has suggested that the increased infectivity may be the result of enhanced functional S protein assembly on the surface of the virion. Several other studies have reported that D614G may be associated with higher viral loads. Fortunately, since this mutation became common to nearly all sequenced SARS-CoV-2 genomes before the release of COVID-19 vaccines, we can be confident that vaccines with proven efficacy against SARS-CoV-2 are protective against the D614G mutation.

The N-terminal S1 subunit of the S protein is responsible for the virus-receptor binding of SARS-CoV-2. Research indicates that the acquisition of nucleotide deletions in the amino (N)-terminal domain (NTD) of the S protein may alter antigenicity. According to the Centers for Disease Control and Prevention (CDC), the deletion of amino acids 69 and 70 in B.1.1.7, is likely to cause a conformational change in the spike protein. And the creation of a Δ69Δ70 deletion mutant via site-directed mutagenesis and lentiviral pseudo typing resulted in 2-fold higher infectivity than the WT (D614G background), indicating that this linked pair of amino acid deletions may improve SARS-CoV-2 fitness. Deletion of amino acid 144 in B.1.1.7 and amino acids 242-244 in B.1.351 have also been associated with a reduced binding capacity of certain neutralizing antibodies. Substitution of aspartate with glycine at position 253 (D253G), a mutation that appears in one of the 2 identified forms of B.1.526, has been correlated with an escape from monoclonal antibodies against the NTD, as have L18F, a leucine (L) to phenylalanine (F) substitution at position 18 in P.1 and R246I, an arginine (R) to isoleucine (I) substitution at position 246 in B.1.351. B.1.351, P.1, B.1.427/B.1.429. and B.1.526 all have additional amino acid substitutions in the NTD that are still of unknown significance.

The receptor binding domain (RBD) of the S protein is comprised of amino acids 319-541. It binds directly to ACE-2 receptors in human cells. Therefore, mutations in this portion of the genome are particularly significant to SARS-CoV-2 fitness and antigenicity. B.1.1.7, B.1.351, and P.1 all have a mutation that replaces asparagine (N) with tyrosine (Y) at position 501 of the RBD. N501Y has been shown to increase the binding capacity of SARS-CoV-2 to human ACE-2 receptors, disrupt antibody binding to RBD, and has been implicated in reduced antibody production via weakened T and B cell cooperation. Together, these findings suggest that SARS-CoV-2 variants possessing the N501Y mutation may have an increased potential for immune escape. B.1.351 and P.1 have 2 additional RBD mutations in common, K417N or K417T, a lysine (K) to asparagine (N) or threonine (T) substitution at position 417, and E484K, a glutamate (E) to lysine (K) substitution at position 484. E484K increases the affinity of RBD for ACE-2, increases resistance to SARS-CoV-2 neutralizing antibodies, is less responsive to monoclonal antibody therapy, and reduces neutralization against convalescent plasma. Studies have demonstrated that, in combination, these 3 RBD mutations induce a relatively high conformational change, compared to N501Y alone or the WT strain, indicating the increased potential for immune escape. As mentioned above, B.1.526, has been detected by West Jr. et al. in 2 forms. One of these contains E484K, while the other contains S477N, a serine (S) to asparagine (N) substitution at position 477 that has also been shown to increase receptor binding affinity, suggesting that both forms may demonstrate increased viral infectivity. The variant introduced by Zhang et al. as CAL.20C has also been detected in two forms, B.1.427 and B.1.429, both of which contain the same 3 S gene mutations that are not present in B.1.1.7, B.1.351, P.1 or B.1.526. One of these, L452R, is a substitution that replaces leucine (L) with arginine (R) at position 452 of the RBD and increases the affinity of RBD for ACE-2. Reports of a study conducted by Chui et al. at USCF, indicate that B.1.429 is less susceptible to neutralizing antibodies and may be linked to worse outcomes of disease.

The furin cleavage site of S protein subunits S1 and S2 is essential for the membrane fusion of SARS-CoV-2. Loss of this structure or function has a major negative impact on the pathogenesis of the virus. B.1.1.7 has a proline (P) to histidine (H) substitution at position 681 which is located near the furin cleavage site. This mutation may further impact viral infectivity, although it is not yet clear if P681H enhances or decreases infectivity. B.1.351 and B.1.526 both have an alanine (A) to valine (V) substitution located adjacent to the furin cleavage site at position 701 (A701V) that is still of unknown significance. Hodcroft et al., have identified a rapid rise of SARS-CoV-2 infections that possess a substitution at position 677 of the S gene (34). It was suspected that the proximity of the mutation to the furin cleavage site may impact the virus’ ability to enter host cells, and parallel evolution in multiple lineages could suggest a selective advantage to the virus. So far, one sub-lineage carrying Q677P, glutamine (Q) to proline (P) substitution, and at least 6 distinct sub-lineages carrying Q677H, glutamine (Q) to histidine (H) substitution have been detected, demonstrating the importance of continued research to evaluate these S:677 polymorphisms.

In conclusion, we report the first universal Coronavirus vaccine was safe, immunogenic, and provided cross-protection against six SARS-CoV-2 variants of Concern.

## MATERIALS & METHODS

### Viruses

SARS-CoV-2 viruses specific to six variants, namely (*i*) SARS-CoV-2-USA/WA/2020 (Batch Number: G2027B); (*ii*) Alpha (B.1.1.7) (isolate England/204820464/2020 Batch Number: C2108K); (*iii*) Beta (B.1.351) (isolate South Africa/KRISP-EC-K005321/2020; Batch Number: C2108F), (*iv*) Gamma (P.1) (isolate hCoV-19/Japan/TY7-503/2021; Batch Number: G2126A), (v) Delta (B.1.617.2) (isolate h-CoV-19/USA/MA29189; Batch number: G87167), and Omicron (BA.1.529) (isolate h-CoV-19/USA/FL17829; Batch number: G76172) were procured from Microbiologics (St. Cloud, MN). The initial batches of viral stocks were propagated to generate high-titer virus stocks. Vero E6 (ATCC-CRL1586) cells were used for this purpose using an earlier published protocol (35). Procedures were completed only after appropriate safety training was obtained using an aseptic technique under BSL-3 containment.

### Triple transgenic mice immunization with SARS-CoV-2 conserved peptides and Infection

The University of California-Irvine conformed to the Guide for the Care and Use of Laboratory Animals published by the US National Institute of Health (IACUC protocol # AUP-22-086). Seven to eight-week-old triple transgenic HLA-A*02:01/HLA-DRB1*01:01-hACE-2 mice (n=60) were included in this experiment. Mice were subcutaneously immunized with a pool of conserved Pan-Coronavirus peptides. The peptide pool administered per mouse comprised 25_μ_g each of the 9-mer long 16 CD8^+^ T cell peptides (ORF1ab_2210-2218_, ORF1ab_3013-3021_, ORF1ab_4283-4291_, ORF1ab_6749-6757_, ORF6_3-11_, ORF7b_26-34_, ORF8a_73-81_, ORF10_3-11_, ORF10_5-13_, S_958-966_, S_1000-1008_, S_1220-1228_, E_20-28_, E_26-34_, M_52-60_, and M_89-97_), 15-mer long 6 CD4^+^ T cell epitopes (ORF1a_1350-1365_, ORF6_12-26_, ORF8b_1-15_, S_1-13_, M_176-190_, and N_388-403_), and 9 B-cell peptides. The pool of peptides was then mixed with 25_μ_g of CpG and 25_μ_g of Alum to prepare the final composition. Mice were immunized with the peptide pool on Day 0 and Day 14 of the experiment. Fourteen days following the second immunization, on Day 28, mice were divided into 6 groups and intranasally infected with 1 × 10^5^ pfu of SARS-CoV-2 (USA-WA1/2020) (n=10), 6 × 10^3^ pfu of SARS-CoV-2-Alpha (B.1.1.7) (n=10), 6 × 10^3^ pfu of SARS-CoV-2-Beta (B.1.351) (n=10), 5 × 10^2^ pfu of SARS-CoV-2-Gamma (P.1) (n=10), 8 × 10^3^ pfu of SARS-CoV-2-Delta (B.1.617.2) (n=10), and 6.9 × 10^4^ pfu of SARS-CoV-2-Omicron (B.1.1.529) (n=10). The viruses were diluted, and each mouse was administered intranasally with 20_μ_l volume. Mice were monitored daily for weight loss and survival until Day 14 p.i. Throat swabs were collected for viral titration on Days 2, 4, 6, 8, 10, and 14 post-infection.

### Human study population cohort and HLA genotyping

In this study, we have included 210 subjects from a pool of over 682 subjects. Written informed consent was obtained from participants before inclusion. The subjects were categorized as mild to severe COVID-19 groups and have undergone treatment at the University of California Irvine Medical Center between July 2020 to July 2022 (Institutional Review Board protocol #-2020-5779). SARS-CoV-2 positivity was defined by a positive RT-PCR on nasopharyngeal swab samples. All the subjects were genotyped by PCR for class I HLA-A*02:01 and class II HLA-DRB1*01:01 among the 682 patients (and after excluding a few for which the given amount of blood was insufficient – i.e., less than 6ml), we ended up with 210 that were genotyped for HLA-A*02:01^+^ or/and HLA-DRB1*01:01^+^ (^36, 37^). Based on the severity of symptoms and ICU admission/intubation status, the subjects were divided into five broad severity categories namely: Severity 5: patients who died from COVID-19 complications; Severity 4: infected COVID-19 patients with severe disease that were admitted to the intensive care unit (ICU) and required ventilation support; Severity 3: infected COVID-19 patients with severe disease that required enrollment in ICU, but without ventilation support; Severity 2: infected COVID-19 patients with moderate symptoms that involved a regular hospital admission; Severity 1: infected COVID-19 patients with mild symptoms; and Severity 0: infected individuals with no symptoms. Demographically, the 210 patients included were from mixed ethnicities (Hispanic (34%), Hispanic Latino (29%), Asian (19%), Caucasian (14%), Afro-American (3%), and Native Hawaiian and Other Pacific Islander descent (1%).

### Sequence comparison among variants of SARS-CoV-2 and animal CoV strains

We retrieved nearly 8.5 million human SARS-CoV-2 genome sequences from the GISAID database representing countries from North America, South America, Central America, Europe, Asia, Oceania, Australia, and Africa. This comprised all the VOCs and VBMs of SARS-CoV-2 (B.1.177, B.1.160, B.1.1.7, B.1.351, P.1, B.1.427/B.1.429, B.1.258, B.1.221, B.1.367, B.1.1.277, B.1.1.302, B.1.525, B.1.526, S:677H.Robin1, S:677P.Pelican, B.1.617.1, B.1.617.2, B,1,1,529) and common cold SARS-CoV strains (SARS-CoV-2-Wuhan-Hu-1 (MN908947.3), SARS-CoV-Urbani (AY278741.1), HKU1-Genotype B (AY884001), CoV-OC43 (KF923903), CoV-NL63 (NC_005831), CoV-229E (KY983587)) and MERS (NC_019843)). Also, for evaluating the evolutionary relationship among the SARS-CoV-2 variants and common cold CoV strains, we have included whole-genome sequences from the bat ((RATG13 (MN996532.2), ZXC21 (MG772934.1), YN01 (EPI_ISL_412976), YN02(EPI_ISL_412977), WIV16 (KT444582.1), WIV1 (KF367457.1), YNLF_31C (KP886808.1), Rs672 (FJ588686.1)), pangolin (GX-P2V (MT072864.1), GX-P5E (MT040336.1), GX-P5L (MT040335.1), GX-P1E (MT040334.1), GX-P4L (MT040333.1), GX-P3B (MT072865.1), MP789 (MT121216.1), Guangdong-P2S (EPI_ISL_410544)), camel (KT368891.1, MN514967.1, KF917527.1, NC_028752.1), and civet (Civet007, A022, B039)). All the sequences included in this study were retrieved either from the NCBI GenBank (www.ncbi.nlm.nih.gov/nuccore) or GISAID (www.gisaid.org). Multiple sequence alignment was performed keeping SARS-CoV-2-Wuhan-Hu-1 (MN908947.3) protein sequence as a reference against all the SARS-CoV-2 VOCs, common cold, and animal CoV strains. The sequences were aligned using the ClustalW algorithm in DIAMOND (38).

### SARS-CoV-2 CD8^+^ and CD4^+^ T Cell Epitope Prediction

Epitope prediction was performed considering the spike glycoprotein (YP_009724390.1) for the reference SARS-CoV-2 isolate, Omicron BA.2. The reference spike protein sequence was used to screen CD8^+^ T cell and CD4^+^ T cell epitopes. The tools used for CD8+ T cell-based epitope prediction were SYFPEITHI, MHC-I binding predictions, and Class I Immunogenicity. Of these, the latter two were hosted on the IEDB platform. We used multiple databases and algorithms for the prediction of CD4^+^ T cell epitopes, namely SYFPEITHI, MHC-II Binding Predictions, Tepitool, and TEPITOPEpan. For CD8^+^ T cell epitope prediction, we selected the 5 most frequent HLA-A class I alleles (HLA-A*01:01, HLA-A*02:01, HLA-A*03:01, HLA-A*11:01, HLA-A*23:01) with nearly 91.48% coverage of the world population, regardless of race and ethnicity, using a phenotypic frequency cutoff ≥ 6%. Similarly, for CD4^+^ T cell epitope prediction, selected HLA-DRB1*01:01, HLA-DRB1*11:01, HLA-DRB1*15:01, HLA-DRB1*03:01, HLA-DRB1*04:01 alleles with population coverage of 86.39%. Subsequently, using NetMHC we analyzed the SARS-CoV-2 protein sequence against all the MHC-I and MHC-II alleles. Epitopes with 9-mer lengths for MHC-I and 15-mer lengths for MHC-II were predicted. Subsequently, the peptides were analyzed for binding stability to the respective HLA allotype. Our stringent epitope selection criteria were based on picking the top 1% epitopes focused on prediction percentile scores. N and O glycosylation sites were screened using NetNGlyc 1.0 and NetOGlyc 4.0 prediction servers, respectively.

### Population-Coverage-Based T Cell Epitope Selection

For a robust epitope screening, we evaluated the conservancy of CD8^+^ T cell, CD4^+^ T cell, and B cell epitopes within spike glycoprotein of Human-SARS-CoV-2 genome sequences representing North America, South America, Africa, Europe, Asia, and Australia. As of April 20th, 2022, the GISAID database extrapolated 8,559,210 human-SARS-CoV-2 genome sequences representing six continents. Population coverage calculation (PPC) was carried out using the Population Coverage software hosted on the IEDB platform. PPC was performed to evaluate the distribution of screened CD8^+^ and CD4^+^ T cell epitopes in the world population at large in combination with HLA-I (HLA-A*01:01, HLA-A*02:01, HLA-A*03:01, HLA-A*11:01, HLA-A*23:01), and HLA-II (HLA-DRB1*01:01, HLA-DRB1*11:01, HLA-DRB1*15:01, HLA-DRB1*03:01, HLA-DRB1*04:01) alleles.

### T cell epitopes screening, selection, and peptide synthesis

Peptide-epitopes from twelve SARS-CoV-2 proteins, including 9-mer long 16 CD8^+^ T cell epitopes (ORF1ab_2210-2218_, ORF1ab_3013-3021_, ORF1ab_4283-4291_, ORF1ab_6749-6757_, ORF6_3-11_, ORF7b_26-34_, ORF8a_73-81_, ORF10_3-11_, ORF10_5-13_, S_958-966_, S_1000-1008_, S_1220-1228_, E_20-28_, E_26-34_, M_52-60_, and M_89-97_) and 15-mer long 6 CD4^+^ T cell epitopes (ORF1a_1350-1365_, ORF6_12-26_, ORF8b_1-15_, S_1-13_, M_176-190_, and N_388-403_) that we formerly identified were selected as described previously. (33) The Epitope Conservancy Analysis tool was used to compute the degree of identity of CD8^+^ T cell and CD4^+^ T cell epitopes within a given protein sequence of SARS-CoV-2 set at 100% identity level (33). Peptides were synthesized as previously described (21^st^ Century Biochemicals, Inc, Marlborough, MA). The purity of peptides determined by both reversed-phase high-performance liquid chromatography and mass spectroscopy was over 95%. Peptides were first diluted in DMSO and later in PBS (1 mg/mL concentration). The helper T-lymphocyte (HTL) epitopes for the selected SARS-CoV-2 proteins were predicted using the MHC-II epitope prediction tool from the Immune Epitope Database (IEDB, http://tools.iedb.org/mhcii/). Selected epitopes had the lowest percentile rank and IC_50_ values. Additionally, the selected epitopes were checked by the IFN epitope server (http://crdd.osdd.net/raghava/ifnepitope/) for the capability to induce Th1 type immune response accompanied by IFN-[production. Cytotoxic T-lymphocyte (CTL) epitopes for the screened proteins were predicted using the NetCTL1.2 server (http://www.cbs.dtu.dk/services/NetCTL/). B-cell epitopes for the screened SARS-CoV-2 proteins were predicted using the ABCPredserver (http://crdd.osdd.net/raghava/abcpred/). The prediction of the toxic/non-toxic nature of all the selected HTL, CTL, and B-cell epitopes was checked using the ToxinPred module(http://crdd.osdd.net/raghava/toxinpred/multi_submit.php).

### Immunogenicity and allergenicity prediction

The immunogenicity of the vaccine was determined using the VaxiJen server (http://www.ddg-pharmfac.net/vaxijen/VaxiJen/VaxiJen.html) and ANTIGEN pro module of SCRATCH protein predictor (http://scratch.proteomics.ics.uci.edu/). The allergenicity of the vaccine was checked using AllerTOPv2.0 (http://www.ddg-pharmfac.net/AllerTOP/) and AlgPredServer (http://crdd.osdd.net/raghava/algpred/).

### SARS-CoV-2 B Cell Epitope Prediction

Linear B cell epitope predictions were carried out on the spike glycoprotein (S), the primary target of B cell immune responses for SARS-CoV. We used the BepiPred 2.0 algorithm embedded in the B cell prediction analysis tool hosted on the IEDB platform. For each protein, the epitope probability score for each amino acid and the probability of exposure was retrieved. Potential B cell epitopes were predicted using a cutoff of 0.55 (corresponding to a specificity greater than 0.81 and sensitivity below 0.3) and considering sequences having more than 5 amino acid residues. This screening process resulted in 8 B-cell peptides. These epitopes represent all the major non-synonymous mutations reported among the SARS-CoV-2 variants. One B-cell epitope (S_439-482_) was observed to possess the maximum number of variant-specific mutations. Structure-based antibody prediction was performed using Discotope 2.0, and a positivity cutoff greater than −2.5 was applied (corresponding to specificity greater than or equal to 0.80 and sensitivity below 0.39), using the SARS-CoV-2 spike glycoprotein structure (PDB ID: 6M1D).

### Determination of physicochemical properties

The physiochemical characteristics of the vaccine were determined using the ProtParam tool of the ExPASy database server (http://web.expasy.org/protparam/).

### Structure prediction, validation, and docking with the receptor

The secondary structure of the subunit, the vaccine construct was predicted using PSIPred4.0 Protein Sequence Analysis Workbench(http://bioinf.cs.ucl.ac.uk/psipred/), while the tertiary structure was predicted by de novo structure prediction based trRosetta modeling suite, which uses a deep residual neural network to predict the inter-residue distance and orientation distribution of the input sequence. Then it converts predicted distance and orientation distribution into smooth restraints to build a 3D structure model based on direct energy minimization. The model of the vaccine construct with the best TM-score was validated by PROCHECKv3.5 (https://servicesn.mbi.ucla.edu/PROCHECK/) and ProSA(https://prosa.services.came.sbg.ac.at/prosa.php) web servers.

### Molecular dynamics simulations

Molecular dynamics (MD) simulation is an effective method to study the molecular interactions and dynamics of the vaccine-ACE-2 complex. The complex structure of the vaccine was initially optimized using Schrödinger Maestro (Schrödinger Release 2016–4: Maestro, Schrödinger, New York) and subsequently used as the starting structure for MD simulations. First, hydrogen atoms were added to the complex which was then solvated in an octahedral box with a simple point charge (SPC) water in the center at least 1.0nm from the box edge. The system was subsequently electrostatically neutralized by the addition of appropriate counter ions. MD simulation was carried out with GROMACS 5.1.2 software package using the gromos9654A7 force field. A standard MD simulation protocol started with 50,000 steps of energy minimization until no notable change of energy was observed, followed by a heating step from 0 to 300K in 200ps (canonical ensemble) and 1000 psat300K (isobaric isothermal ensemble) by constant temperature equilibration. During this, Parrinello-Rahman barostat pressure coupling was used to avoid the impact of velocity. As a final step of the simulation, a 40ns production run was carried out at 300K with periodic boundary conditions in the NPT ensemble with modified Bendensen temperature coupling and at a constant pressure of 1 atm. Furthermore, the LINCSalgorithm, along with the Particlemest Ewald method, was used for the calculation of the long-range electrostatic forces. Fourier grid pacing and Coulomb radius were set at 0.16 and 1.4 nm respectively, during the simulations. The van der Waals (VDW) interactions were limited to 1.4nm, and structures were saved at every 10ps for structural and dynamic analysis.

### Sequence-based variant effect prediction

Fourteen spike glycoprotein-specific non-synonymous mutations found on different SARS-CoV-2 variants were subjected to Variant Effect Predictor (VEP) online server for effect prediction against SARS-CoV-2 genome assembly hosted by Ensemble database. VEP was set to return (i) Combined Annotation Dependent Depletion (CADD) score, (ii) Genomic Evolutionary Rate Profiling (GERP ++ Raw Score), (iii) phastCons conservation score based on the multiple alignments of 7 vertebrate genomes, (iv) phylogenetic p-values (PhyloP) conservation score based on the multiple alignments of 7 vertebrate genomes, (v) Shifting Intolerant From Tolerant (SIFT) score and prediction, (vi) Polymorphism Phenotyping (PolyPhen) score and prediction, (vii) Consensus Deleteriousness (Condel) rank score and prediction, (viii) Protein Variation Effect Analyzer (PROVEAN) score and prediction, and (ix) Mutation Accessor score and prediction.

### Blood Differential Test (BDT)

Total White Blood Cells (WBCs) count and Lymphocytes count per µL of blood were performed by the University of California Irvine Medical Center clinicians using CellaVision^TM^ DM96 automated microscope. Monolayer smears were prepared from anticoagulated blood and stained using the May Grunwald Giemsa (MGG) technique. Subsequently, slides were loaded onto the DM96 magazines and scanned using a 10-x objective focused on nucleated cells to record their exact position. Images were obtained using the 100-x oil objective and analyzed by Artificial Neural Network (ANN).

### TaqMan quantitative polymerase reaction assay for the screening of SARS-CoV-2 Variants in COVID-19 patients

We utilized a laboratory-developed modification of the CDC SARS-CoV-2 RT-PCR assay, which received Emergency Use Authorization by the FDA on 17 April 2020. (https://www.fda.gov/media/137424/download [accessed 24 March 2021]).

Mutation screening assays: SARS-CoV-2-positive samples were screened by four multiplex RT-PCR assays. Through the qRT-PCR, we screened for 11 variants of SARS-CoV-2 in our patient cohort. The variants which were screened include B.1.1.7 (Alpha), B.1.351 (Beta), P.1 (Gamma), and B.1.427/B.1.429 (Epsilon), B.1.525 (Eta), R.1, P.2 (Zeta), B.1.526 (Iota), B.1.2/501Y or B.1.1.165, B.1.1.529 (BA.1) (Omicron), B.1.1.529 (BA.2) (Omicron), and B.1.617.2 (Delta). The sequences for the detection of Δ69–70 were adapted from a multiplex real-time RT-PCR assay for the detection of SARS-CoV-2 (Zhen et al., 2020). The probe overlaps with the sequences that contain amino acids 69 to 70; therefore, a negative result for this assay predicts the presence of deletion S-Δ69–70 in the sample. Using a similar strategy, a primer/probe set that targets the deletion S-Δ242–244 was designed and was run in the same reaction with S-Δ69-70. In addition, three separate assays were designed to detect spike mutations S-501Y, S-484K, and S-452R and wild-type positions S-501N, S-484E, and S-452L.

Briefly, 5 ml of the total nucleic acid eluate was added to a 20-*m*l total-volume reaction mixture (1x TaqPath 1-Step RT-qPCR Master Mix, CG [Thermo Fisher Scientific, Waltham, MA], with 0.9 *m*M each primer and 0.2 *m*M each probe). The RT-PCR was carried out using the ABI StepOnePlus thermocycler (Life Technologies, Grand Island, NY). The S-N501Y, S-E484K, and S-L452R assays were carried out under the following running conditions: 25°C for 2 min, then 50°C for 15 min, followed by 10 min at 95°C and 45 cycles of 95°C for 15 s and 65°C for 1 min. The Δ 69–70 / Δ242–244 assays were run under the following conditions: 25°C for 2 min, then 50°C for 15 min, followed by 10 min at 95°C and 45 cycles of 95°C for 15 s and 60°C for 1 min. Samples displaying typical amplification curves above the threshold were considered positive. Samples that yielded a negative result or results in the S-Δ69–70/ Δ242–244 assays or were positive for S-501Y P2, S-484K P2, and S-452R P2 were considered screen positive and assigned to a VOC.

### Neutralizing antibody assays for SARS-CoV-2

Serially diluted heat-inactivated plasma (1:3) and 300 PFU of SARS-CoV-2 variants are combined in Dulbecco’s Modified Eagle’s Medium (DMEM) and incubated at 37°C 5% CO2 for 30 minutes. After neutralization, the antibody-virus inoculum was transferred onto Vero E6 cells (ATCC C1008) and incubated at 34°C 5% CO2 for 1 hour. The Vero cells were seeded in a 96-well plate at 3.5×10^4^ cells/well 24 hours before the assay. After 1 hour, 1% methylcellulose (Sigma Aldrich) at a 1:1 ratio was overlaid on the infected Vero cell layer. Plates were incubated at 34°C 5% CO_2_ for 24 hours. After 24 hours, the medium was carefully removed, and the plates were fixed with 100μl of 10% neutral buffered formalin for 1 hour at room temperature. Following fixation, plates were washed 3 times using deionized (DI) water, and 50μl of ice-cold Methanol supplemented with 0.3% hydrogen peroxide was added to each well. Plates were incubated at −20°C for 10 minutes followed by 20 minutes at room temperature. Methanol/Hydrogen peroxide was removed by washing the plates 3 times with DI water. Once washed, plates were blocked for 1 hour with 5% non-fat dry milk in PBS. The blocking solution was removed and 40μl/well of anti-SARS Nucleocapsid antibody (Novus Biologicals NB100-56576) at 1:1000 in 5% non-fat dry milk/PBS was added. Plates were incubated overnight at 4°C followed by a 2-hour room temperature incubation (with agitation). Plates were washed 4 times with PBS and 40μl of HRP anti-rabbit IgG antibody (Biolegend) 1:1500 was added to each well and incubated for 2 hours at room temperature. Plates were developed using True Blue HRP substrate and imaged on an ELIPOT reader. Each plate was set up with a positive neutralization control and a negative control (Virus/no plasma). The half maximum inhibitory concentration (IC50) was calculated by non-linear regression analysis using normalized counted foci on Prism 7 (GraphPad Software). 100% infectivity was obtained by normalizing the number of foci counted in the wells derived from the cells infected with the SARS-CoV-2 virus in the absence of plasma.

### Histology of animal lungs

Mouse lungs were preserved in 10% neutral buffered formalin for 48 hours before transferring to 70% ethanol. The tissue sections were then embedded in paraffin blocks and sectioned at 8 μm thickness. Slides were deparaffinized and rehydrated before staining for hematoxylin and eosin for routine immunopathology. IHC was performed on mice lung tissues probed with SARS/SARS-CoV-2 Coronavirus NP Monoclonal Antibody (B46F) (Product # MA1-7404) at a dilution of 1:100. The antibody showed significant staining in lung tissues of non-immunized, SARS-CoV-2 infected mice when compared to the tissues of the vaccinated group of mice. This method was meant to demonstrate the relative expression of the Nucleocapsid protein between non-immunized Mock and immunized samples. Further CD8^+^ T cell and CD4^+^ T cell-specific staining were performed to identify the T cell infiltration among the immunized and Mock groups.

### Peripheral blood mononuclear cells isolation and T cell stimulation

Peripheral blood mononuclear cells (PBMCs) from COVID-19 patients were isolated from the blood using Ficoll (GE Healthcare) density gradient media and transferred into 96-well plates at a concentration of 2.5 × 10^6^ viable cells per ml in 200µl (0.5 × 10^6^ cells per well) of RPMI-1640 media (Hyclone) supplemented with 10% (v/v) FBS (HyClone), Sodium Pyruvate (Lonza), L-Glutamine, Nonessential Amino Acids, and antibiotics (Corning). A fraction of the blood was kept separated to perform HLA genotyping of the patients and select only the HLA-A*02:01 and/or DRB1*01:01 positive individuals. Subsequently, cells were then stimulated with 10µg/ml of each one of the 22 individual T cell peptide-epitopes (16 CD8^+^ T cell peptides and 6 CD4^+^ T cell peptides) and incubated in humidified 5% CO_2_ at 37°C. Post-incubation, cells were stained by flow cytometry analysis, or transferred in IFN-γ ELISpot plates. The same isolation protocol was followed for healthy donor (HD) samples obtained in 2018. PBMC samples were kept frozen in liquid nitrogen in 10% FBS in DMSO. Upon thawing, HD PBMCs were stimulated in the same manner for the IFN-γ ELISpot technique.

### ELISpot assay

COVID-19 patients were first screened for their HLA status (DRB1*01:01^+^ positive = 108, HLA-A*02:01^+^ positive = 83, DRB1*01:01^+^ and HLA-A*02:01^+^ positive = 19). The 108 DRB1*01:01 positive individuals were used to assess the CD4^+^ T-cell response against our SL-CoVs-conserved SARS-CoV-2-derived class-II restricted epitopes by IFN-γ ELISpot. Subsequently, we assessed the CD8^+^ T cell response against our SL-CoVs conserved SARS-CoV-2 derived class-I restricted epitopes in the 83 HLA-A*02:01 positive individuals representing different disease severity categories. ELISpot assay was performed as described previously in (33, 39).

### Flow cytometry analysis

After 72 hours of stimulation with each SARS-CoV-2 class-I or class-II restricted peptide, PBMCs (0.5 × 10^6^ cells) from 147 patients were stained for the detection of surface markers and subsequently analyzed by flow cytometry. First, the cells were stained with a live/dead fixable dye (Zombie Red dye, 1/800 dilution – BioLegend, San Diego, CA) for 20 minutes at room temperature, to exclude dying/apoptotic cells. Cells were stained for 45 minutes at room temperature with five different HLA-A*02*01 restricted tetramers and/or five HLA-DRB1*01:01 restricted tetramers (PE-labelled) specific toward the SARS-CoV-2 CD8^+^ T cell epitopes Orf1ab_2210-2218_, Orf1ab_4283-4291_, S_1220-1228_, ORF10_3-11_ and toward the CD4^+^ T cell epitopes ORF1a_1350-1365_, S_1-13_, M_176-190_, ORF6_12-26_, respectively. We have optimized our tetramer staining according to the instructions published by Dolton et al. (40) As a negative control aiming to assess tetramer staining specificity, we stained HLA-A*02*01-HLA-DRB1*01:01-negative patients with our four tetramers. Subsequently, we used anti-human antibodies for surface marker staining: anti-CD62L, anti-CD69, anti-CD4, anti-CD8, and anti-IFN-g. mAbs against these various cell markers were added to the cells in phosphate-buffered saline (PBS) containing 1% FBS and 0.1% sodium azide (fluorescence-activated cell sorter [FACS] buffer) and left for 30 minutes at 4°C. At the end of the incubation period, the cells were washed twice with FACS buffer and fixed with 4% paraformaldehyde (PFA, Affymetrix, Santa Clara, CA). A total of ~200,000 lymphocyte-gated PBMCs (140,000 alive CD45^+^) were acquired by Fortessa X20 (Becton Dickinson, Mountain View, CA) and analyzed using FlowJo software (TreeStar, Ashland, OR).

### Enzyme-linked immunosorbent assay (ELISA)

Serum antibodies specific for epitope peptides and SARS-CoV-2 proteins were detected by ELISA. 96-well plates (Dynex Technologies, Chantilly, VA) were coated with 0.5 μg peptides, 100 ng S or N protein per well at 4°C overnight, respectively, and then washed three times with PBS and blocked with 3% BSA (in 0.1% PBST) for 2 h at 37°C. After blocking, the plates were incubated with serial dilutions of the sera (100 μl/well, in two-fold dilution) for 2 hours at 37°C. The bound serum antibodies were detected with HRP-conjugated goat anti-mouse IgG and chromogenic substrate TMB (ThermoFisher, Waltham, MA). The cut-off for seropositivity was set as the mean value plus three standard deviations (3SD) in HBc-S control sera. The binding of the epitopes to the sera of SARS-CoV-2 infected samples was detected by ELISA using the same procedure, 96-well plates were coated with 0.5 μg peptides and sera were diluted at 1:50. All ELISA studies were performed at least twice.

### Data and Code Availability

The human-specific SARS-CoV-2 complete genome sequences were retrieved from the GISAID database, whereas the SARS-CoV-2 sequences for pangolin (Manis javanica), and bat (Rhinolophus affinis, Rhinolophus malayanus) were retrieved from NCBI. Genome sequences of previous strains of SARS-CoV for humans, bats, civet cats, and camels were retrieved from the NCBI GenBank.

## ACKNOWLEDGMENTS

This work is supported by the Fast-Grant PR12501 from Emergent Ventures, by a Gavin Herbert Eye Institute internal grant, by Public Health Service Research grants AI158060, AI174383, AI150091, AI143348, AI147499, AI143326, AI138764, AI124911, and AI110902 from the National Institutes of Allergy and Infectious Diseases (NIAID) to LBM.

---

**<u>Supplemental Figure S1</u>. Sequence homology analysis to identify the degree of the conservancy of the immunodominant CD8^+^ T cell epitopes among SARS-CoV-2 VOCs: Sequence homology data for the CD8^+^ T cell epitopes is shown.** The 16 epitopes, found to be highly immunodominant against SARS-CoV-2 variants of concern WA/USA2020, Alpha (B.1.1.7), Beta (B.1.351), Gamma (P.1), Delta (B.1.617.2), and Omicron (B.1.1.529) were subjected to the sequence homology analysis.

**<u>Supplemental Figure S2</u>**. **Sequence homology analysis to identify the degree of the conservancy of the immunodominant CD4^+^ T cell epitopes among SARS-CoV-2 variants of concern: Sequence homology data for the CD4^+^ T cell epitopes is shown.** The 6 epitopes, found to be highly immunodominant against SARS-CoV-2 variants of concern WA/USA2020, Alpha (B.1.1.7), Beta (B.1.351), Gamma (P.1), Delta (B.1.617.2), and Omicron (B.1.1.529) were subjected to the sequence homology analysis.

**<u>Supplemental Figure S3</u>. Sequence homology analysis to identify the degree of the conservancy of the immunodominant B cell epitopes among SARS-CoV-2 variants of concern: The sequence homology data for the B cell epitopes are shown.** The 9 epitopes, found to be highly immunodominant against SARS-CoV-VOCs WA/USA2020, Alpha (B.1.1.7), Beta (B.1.351), Gamma (P.1), Delta (B.1.617.2), and Omicron (B.1.1.529) were subjected to the sequence homology analysis.

**<u>Supplemental Figure S4</u>. Pan-Coronavirus vaccine evaluation of immunogenicity based on antibody response against “universal” B-cell epitopes in COVID-19 patients and triple transgenic HLA-A*02:01/HLA-DRB1*01:01-hACE-2 exposed to different SARS-CoV-2 variants of concern:** Bar graphs show the peptide binding IgG level for the 9 “universal” B cell epitopes (A) among COVID-19 patients screened to be infected with SARS-CoV-2 variants of concern Alpha (B.1.1.7), Beta (B.1.351), Epsilon (B.1.427/B.1.429), Delta (B.1.617.2), and Omicron (B.1.1.529) and (B) in vaccinated versus mock-vaccinated triple transgenic HLA-A*02:01/HLA-DRB1*01:01-hACE-2 mice. Bars represent means ± SEM. Data were analyzed by student’s *t*-test and multiple t-tests. Results were considered statistically significant at *P* < 0.05. Statistical correction for multiple comparisons was applied using the Holm-Sidak method.

