## Supplemental Figures for "Cross-Protection Induced by Highly Conserved Human B, CD4^+,^ and CD8^+^ T Cell Epitopes-Based Coronavirus Vaccine Against Severe Infection, Disease, and Death Caused by Multiple SARS-CoV-2 Variants of Concern": Prakash Supplemental Figures.pdf


### ABSTRACT

Background: The Coronavirus disease 2019 (COVID-19) pandemic has created one of the largest global health crises in almost a century. Although the current rate of SARS-CoV-2 infections has decreased significantly; the long-term outlook of COVID-19 remains a serious cause of high death worldwide; with the mortality rate still surpassing even the worst mortality rates recorded for the influenza viruses. The continuous emergence of SARS-CoV-2 variants of concern (VOCs), including multiple heavily mutated Omicron sub-variants, have prolonged the COVID-19 pandemic and outlines the urgent need for a next-generation vaccine that will protect from multiple SARS-CoV-2 VOCs. Methods: In the present study, we designed a multi-epitope-based Coronavirus vaccine that incorporated B, $CD4^{+}$, and $CD8^{+}$ T cell epitopes conserved among all known SARS-CoV-2 VOCs and selectively recognized by $CD8^{+}$ and $CD4^{+}$ T-cells from asymptomatic COVID-19 patients irrespective of VOC infection. The safety, immunogenicity, and cross-protective immunity of this pan-Coronavirus vaccine were studied against six VOCs using an innovative triple transgenic h-ACE-2-HLA-A2/DR mouse model. Results: The Pan-Coronavirus vaccine: (*i*) is safe; (*ii*) induces high frequencies of lung-resident functional $CD8^{+}$ and $CD4^{+}$ $T_{EM}$ and $T_{RM}$ cells; and (*iii*) provides robust protection against virus replication and COVID-19-related lung pathology and death caused by six SARS-CoV-2 VOCs: Alpha (B.1.1.7), Beta (B.1.351), Gamma or P1 (B.1.1.28.1), Delta (lineage B.1.617.2) and Omicron (B.1.1.529). Conclusions: A multi-epitope pan-Coronavirus vaccine bearing conserved human B and T cell epitopes from structural and non-structural SARS-CoV-2 antigens induced cross-protective immunity that cleared the virus, and reduced COVID-19-related lung pathology and death caused by multiple SARS-CoV-2 VOCs.

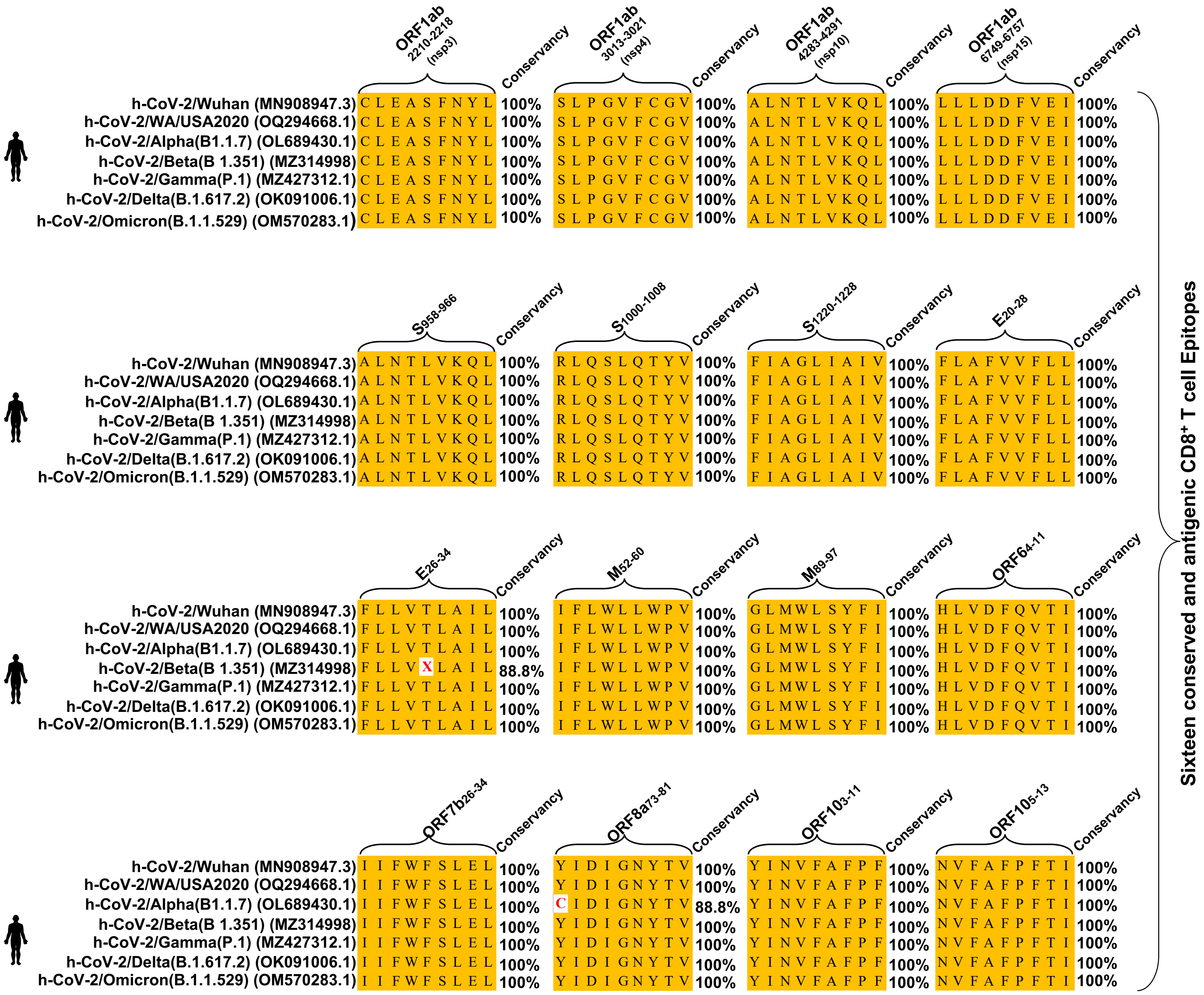
ORF1ab 2210-2218 (nsp3)
Conservancy
ORF1ab 3013-3021 (nsp4)
Conservancy
ORF1ab 4283-4291 (nsp10)
Conservancy
ORF1ab 6749-6757 (nsp15)
Conservancy
h-CoV-2/Wuhan (MN908947.3) C L E A S F N Y L 100% S L P G V F C G V 100% A L N T L V K Q L 100% L L L D D F V E I 100%
h-CoV-2/WA/USA2020 (OQ294668.1) C L E A S F N Y L 100% S L P G V F C G V 100% A L N T L V K Q L 100% L L L D D F V E I 100%
h-CoV-2/Alpha(B1.1.7) (OL689430.1) C L E A S F N Y L 100% S L P G V F C G V 100% A L N T L V K Q L 100% L L L D D F V E I 100%
h-CoV-2/Beta(B 1.351) (MZ314998) C L E A S F N Y L 100% S L P G V F C G V 100% A L N T L V K Q L 100% L L L D D F V E I 100%
h-CoV-2/Gamma(P.1) (MZ427312.1) C L E A S F N Y L 100% S L P G V F C G V 100% A L N T L V K Q L 100% L L L D D F V E I 100%
h-CoV-2/Delta(B.1.617.2) (OK091006.1) C L E A S F N Y L 100% S L P G V F C G V 100% A L N T L V K Q L 100% L L L D D F V E I 100%
h-CoV-2/Omicron(B.1.1.529) (OM570283.1) C L E A S F N Y L 100% S L P G V F C G V 100% A L N T L V K Q L 100% L L L D D F V E I 100%
S958-966
Conservancy
S1000-1008
Conservancy
S1220-1228
Conservancy
E20-28
Conservancy
h-CoV-2/Wuhan (MN908947.3) A L N T L V K Q L 100% R L Q S L Q T Y V 100% F I A G L I A I V 100% F L A F V V F L L 100%
h-CoV-2/WA/USA2020 (OQ294668.1) A L N T L V K Q L 100% R L Q S L Q T Y V 100% F I A G L I A I V 100% F L A F V V F L L 100%
h-CoV-2/Alpha(B1.1.7) (OL689430.1) A L N T L V K Q L 100% R L Q S L Q T Y V 100% F I A G L I A I V 100% F L A F V V F L L 100%
h-CoV-2/Beta(B 1.351) (MZ314998) A L N T L V K Q L 100% R L Q S L Q T Y V 100% F I A G L I A I V 100% F L A F V V F L L 100%
h-CoV-2/Gamma(P.1) (MZ427312.1) A L N T L V K Q L 100% R L Q S L Q T Y V 100% F I A G L I A I V 100% F L A F V V F L L 100%
h-CoV-2/Delta(B.1.617.2) (OK091006.1) A L N T L V K Q L 100% R L Q S L Q T Y V 100% F I A G L I A I V 100% F L A F V V F L L 100%
h-CoV-2/Omicron(B.1.1.529) (OM570283.1) A L N T L V K Q L 100% R L Q S L Q T Y V 100% F I A G L I A I V 100% F L A F V V F L L 100%
E26-34
Conservancy
M52-60
Conservancy
M89-97
Conservancy
ORF64-11
Conservancy
h-CoV-2/Wuhan (MN908947.3) F L L V T L A I L 100% I F L W L L W P V 100% G L M W L S Y F I 100% H L V D F Q V T I 100%
h-CoV-2/WA/USA2020 (OQ294668.1) F L L V T L A I L 100% I F L W L L W P V 100% G L M W L S Y F I 100% H L V D F Q V T I 100%
h-CoV-2/Alpha(B1.1.7) (OL689430.1) F L L V T L A I L 100% I F L W L L W P V 100% G L M W L S Y F I 100% H L V D F Q V T I 100%
h-CoV-2/Beta(B 1.351) (MZ314998) F L L V X L A I L 88.8% I F L W L L W P V 100% G L M W L S Y F I 100% H L V D F Q V T I 100%
h-CoV-2/Gamma(P.1) (MZ427312.1) F L L V T L A I L 100% I F L W L L W P V 100% G L M W L S Y F I 100% H L V D F Q V T I 100%
h-CoV-2/Delta(B.1.617.2) (OK091006.1) F L L V T L A I L 100% I F L W L L W P V 100% G L M W L S Y F I 100% H L V D F Q V T I 100%
h-CoV-2/Omicron(B.1.1.529) (OM570283.1) F L L V T L A I L 100% I F L W L L W P V 100% G L M W L S Y F I 100% H L V D F Q V T I 100%
ORF7b26-34
Conservancy
ORF8a73-81
Conservancy
ORF103-11
Conservancy
ORF105-13
Conservancy
h-CoV-2/Wuhan (MN908947.3) I I F W F S L E L 100% Y I D I G N Y T V 100% Y I N V F A F P F 100% N V F A F P F T I 100%
h-CoV-2/WA/USA2020 (OQ294668.1) I I F W F S L E L 100% Y I D I G N Y T V 100% Y I N V F A F P F 100% N V F A F P F T I 100%
h-CoV-2/Alpha(B1.1.7) (OL689430.1) I I F W F S L E L 100% C I D I G N Y T V 88.8% Y I N V F A F P F 100% N V F A F P F T I 100%
h-CoV-2/Beta(B 1.351) (MZ314998) I I F W F S L E L 100% Y I D I G N Y T V 100% Y I N V F A F P F 100% N V F A F P F T I 100%
h-CoV-2/Gamma(P.1) (MZ427312.1) I I F W F S L E L 100% Y I D I G N Y T V 100% Y I N V F A F P F 100% N V F A F P F T I 100%
h-CoV-2/Delta(B.1.617.2) (OK091006.1) I I F W F S L E L 100% Y I D I G N Y T V 100% Y I N V F A F P F 100% N V F A F P F T I 100%
h-CoV-2/Omicron(B.1.1.529) (OM570283.1) I I F W F S L E L 100% Y I D I G N Y T V 100% Y I N V F A F P F 100% N V F A F P F T I 100%
Sixteen conserved and antigenic CD8+ T cell Epitopes

Prakash et al Supplementary Fig. S2

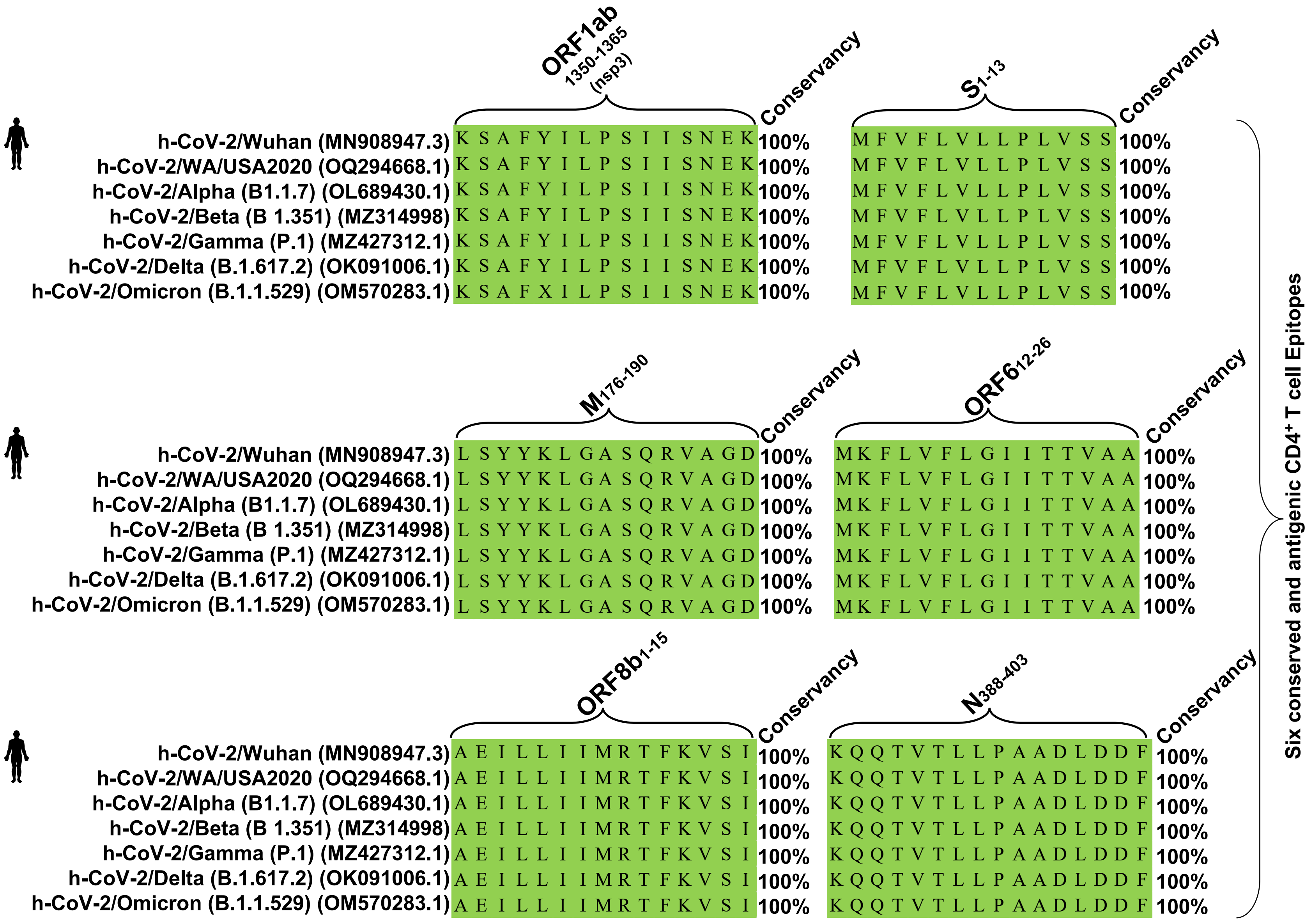

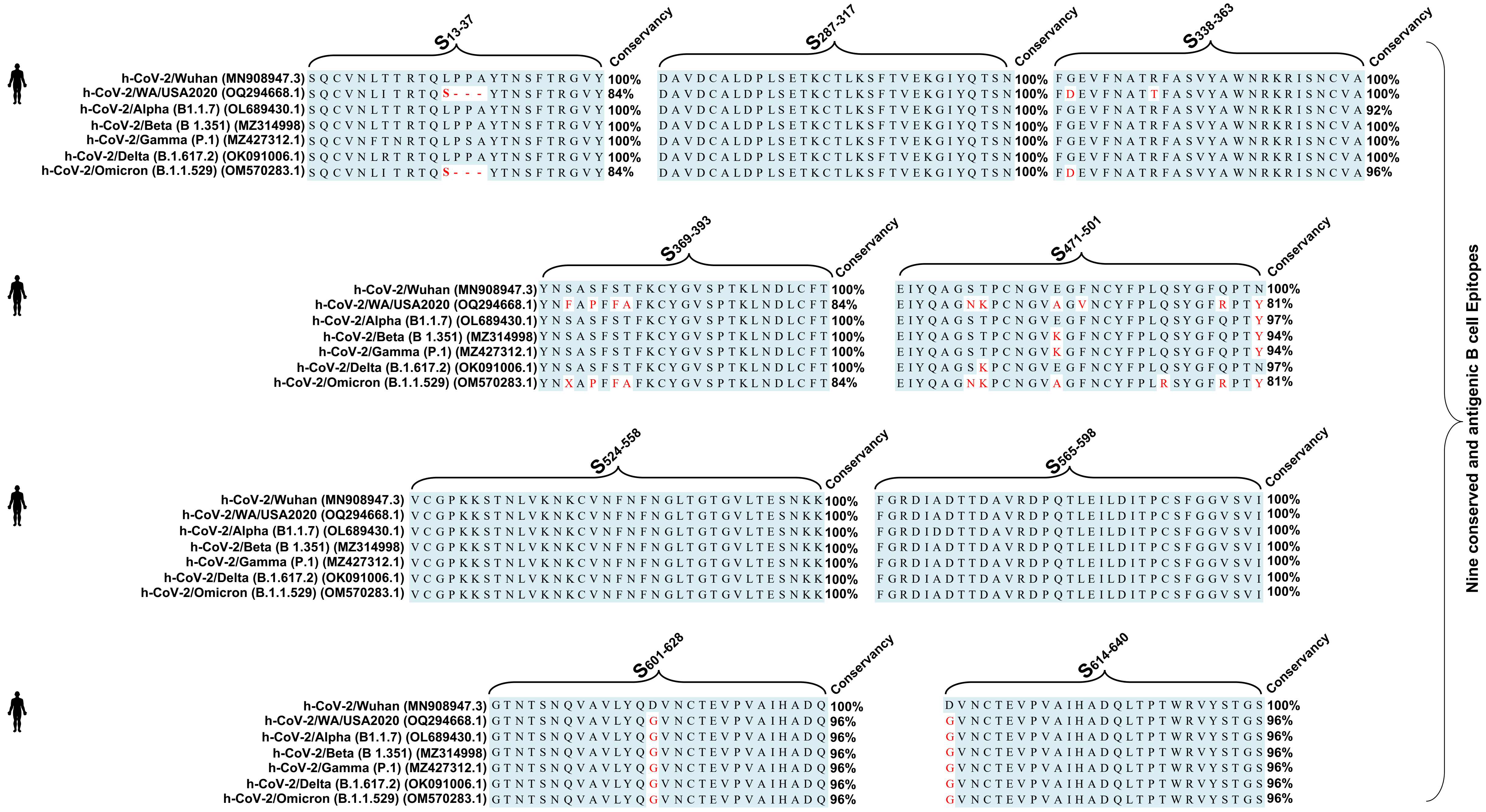
S13-37
Conservancy
h-CoV-2/Wuhan (MN908947.3) SQCVNLTTRTQLPPAYTNSFTRGVY 100%
h-CoV-2/WA/USA2020 (OQ294668.1) SQCVNLI TRTQS---YTNSFTRGVY 84%
h-CoV-2/Alpha (B1.1.7) (OL689430.1) SQCVNLTTRTQLPPAYTNSFTRGVY 100%
h-CoV-2/Beta (B 1.351) (MZ314998) SQCVNLTTRTQLPPAYTNSFTRGVY 100%
h-CoV-2/Gamma (P.1) (MZ427312.1) SQCVNFTNRTQLPSAYTNSFTRGVY 100%
h-CoV-2/Delta (B.1.617.2) (OK091006.1) SQCVNLRTRTQLPPAYTNSFTRGVY 100%
h-CoV-2/Omicron (B.1.1.529) (OM570283.1) SQCVNLI TRTQS---YTNSFTRGVY 84%
S287-317
Conservancy
DAVDCALDPLSETKCTLKSFTVEKGIYQTSN 100%
DAVDCALDPLSETKCTLKSFTVEKGIYQTSN 100%
DAVDCALDPLSETKCTLKSFTVEKGIYQTSN 100%
DAVDCALDPLSETKCTLKSFTVEKGIYQTSN 100%
DAVDCALDPLSETKCTLKSFTVEKGIYQTSN 100%
DAVDCALDPLSETKCTLKSFTVEKGIYQTSN 100%
DAVDCALDPLSETKCTLKSFTVEKGIYQTSN 100%
S338-363
Conservancy
FGEVFNATRFASVYAWNRKRISNCVA 100%
FDEVFNATTFASVYAWNRKRISNCVA 100%
FGEVFNATRFASVYAWNRKRISNCVA 92%
FGEVFNATRFASVYAWNRKRISNCVA 100%
FGEVFNATRFASVYAWNRKRISNCVA 100%
FGEVFNATRFASVYAWNRKRISNCVA 100%
FDEVFNATRFASVYAWNRKRISNCVA 96%
S369-393
Conservancy
h-CoV-2/Wuhan (MN908947.3) YNSASFSTFKCYGVSPTKLNDLCFT 100%
h-CoV-2/WA/USA2020 (OQ294668.1) YNFAPFFAFKCYGVSPTKLNDLCFT 84%
h-CoV-2/Alpha (B1.1.7) (OL689430.1) YNSASFSTFKCYGVSPTKLNDLCFT 100%
h-CoV-2/Beta (B 1.351) (MZ314998) YNSASFSTFKCYGVSPTKLNDLCFT 100%
h-CoV-2/Gamma (P.1) (MZ427312.1) YNSASFSTFKCYGVSPTKLNDLCFT 100%
h-CoV-2/Delta (B.1.617.2) (OK091006.1) YNSASFSTFKCYGVSPTKLNDLCFT 100%
h-CoV-2/Omicron (B.1.1.529) (OM570283.1) YNXAPFFAFKCYGVSPTKLNDLCFT 84%
S471-501
Conservancy
EIYQAGSTPCNGVEGFNCYFPLQSYGFQPTN 100%
EIYQAGNKPCNGVAGVNCYFPLQSYGFRPTY 81%
EIYQAGSTPCNGVEGFNCYFPLQSYGFQPTY 97%
EIYQAGSTPCNGVKGFNCYFPLQSYGFQPTY 94%
EIYQAGSTPCNGVKGFNCYFPLQSYGFQPTY 94%
EIYQAGSKPCNGVEGFNCYFPLQSYGFQPTN 97%
EIYQAGNKPCNGVAGFNCYFPLRSYGFRPTY 81%
S524-558
Conservancy
h-CoV-2/Wuhan (MN908947.3) VCGPKKSTNLVKNKCVNFNFNGLTGTGVLTESNKK 100%
h-CoV-2/WA/USA2020 (OQ294668.1) VCGPKKSTNLVKNKCVNFNFNGLTGTGVLTESNKK 100%
h-CoV-2/Alpha (B1.1.7) (OL689430.1) VCGPKKSTNLVKNKCVNFNFNGLTGTGVLTESNKK 100%
h-CoV-2/Beta (B 1.351) (MZ314998) VCGPKKSTNLVKNKCVNFNFNGLTGTGVLTESNKK 100%
h-CoV-2/Gamma (P.1) (MZ427312.1) VCGPKKSTNLVKNKCVNFNFNGLTGTGVLTESNKK 100%
h-CoV-2/Delta (B.1.617.2) (OK091006.1) VCGPKKSTNLVKNKCVNFNFNGLTGTGVLTESNKK 100%
h-CoV-2/Omicron (B.1.1.529) (OM570283.1) VCGPKKSTNLVKNKCVNFNFNGLTGTGVLTESNKK 100%
S565-598
Conservancy
FGRDIADTTDAVRDPQTLEILDITPCSFGGVSVI 100%
FGRDIADTTDAVRDPQTLEILDITPCSFGGVSVI 100%
FGRDIDDTTDAVRDPQTLEILDITPCSFGGVSVI 100%
FGRDIADTTDAVRDPQTLEILDITPCSFGGVSVI 100%
FGRDIADTTDAVRDPQTLEILDITPCSFGGVSVI 100%
FGRDIADTTDAVRDPQTLEILDITPCSFGGVSVI 100%
FGRDIADTTDAVRDPQTLEILDITPCSFGGVSVI 100%
S601-628
Conservancy
h-CoV-2/Wuhan (MN908947.3) GTNTSNQVAVLYQDVNCTEVPVAIHADQ 100%
h-CoV-2/WA/USA2020 (OQ294668.1) GTNTSNQVAVLYQGVNCTEVPVAIHADQ 96%
h-CoV-2/Alpha (B1.1.7) (OL689430.1) GTNTSNQVAVLYQGVNCTEVPVAIHADQ 96%
h-CoV-2/Beta (B 1.351) (MZ314998) GTNTSNQVAVLYQGVNCTEVPVAIHADQ 96%
h-CoV-2/Gamma (P.1) (MZ427312.1) GTNTSNQVAVLYQGVNCTEVPVAIHADQ 96%
h-CoV-2/Delta (B.1.617.2) (OK091006.1) GTNTSNQVAVLYQGVNCTEVPVAIHADQ 96%
h-CoV-2/Omicron (B.1.1.529) (OM570283.1) GTNTSNQVAVLYQGVNCTEVPVAIHADQ 96%
S614-640
Conservancy
DVNCTEVPVAIHADQLTPTWRVYSTGS 100%
GVNCTEVPVAIHADQLTPTWRVYSTGS 96%
GVNCTEVPVAIHADQLTPTWRVYSTGS 96%
GVNCTEVPVAIHADQLTPTWRVYSTGS 96%
GVNCTEVPVAIHADQLTPTWRVYSTGS 96%
GVNCTEVPVAIHADQLTPTWRVYSTGS 96%
GVNCTEVPVAIHADQLTPTWRVYSTGS 96%
Nine conserved and antigenic B cell Epitopes

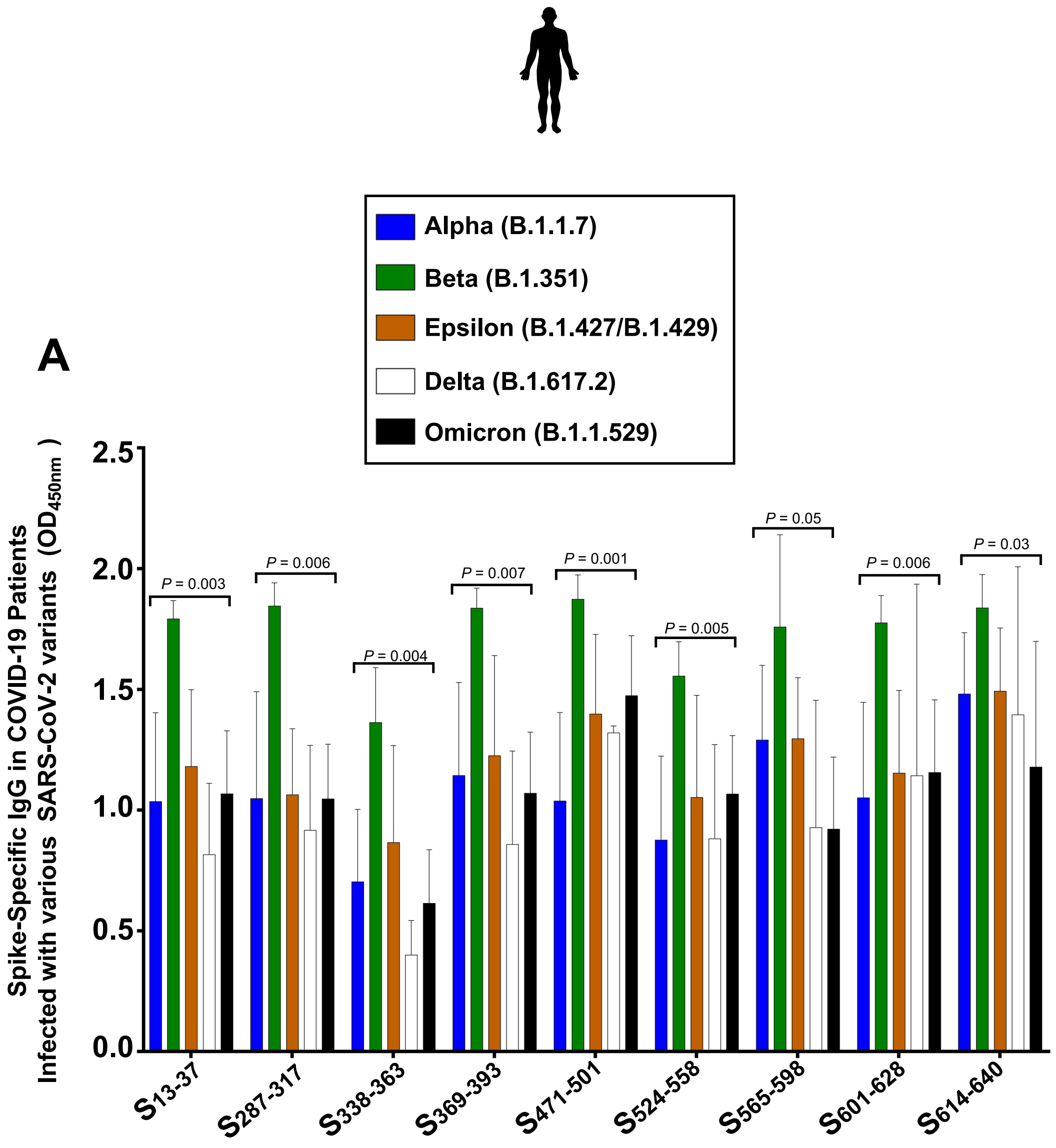
A
Alpha (B.1.1.7)
Beta (B.1.351)
Epsilon (B.1.427/B.1.429)
Delta (B.1.617.2)
Omicron (B.1.1.529)
Spike-Specific IgG in COVID-19 Patients Infected with various SARS-CoV-2 variants (OD450nm)
P = 0.003
P = 0.006
P = 0.004
P = 0.007
P = 0.001
P = 0.005
P = 0.05
P = 0.006
P = 0.03
S13-37
S287-317
S338-363
S369-393
S471-501
S524-558
S565-598
S601-628
S614-640

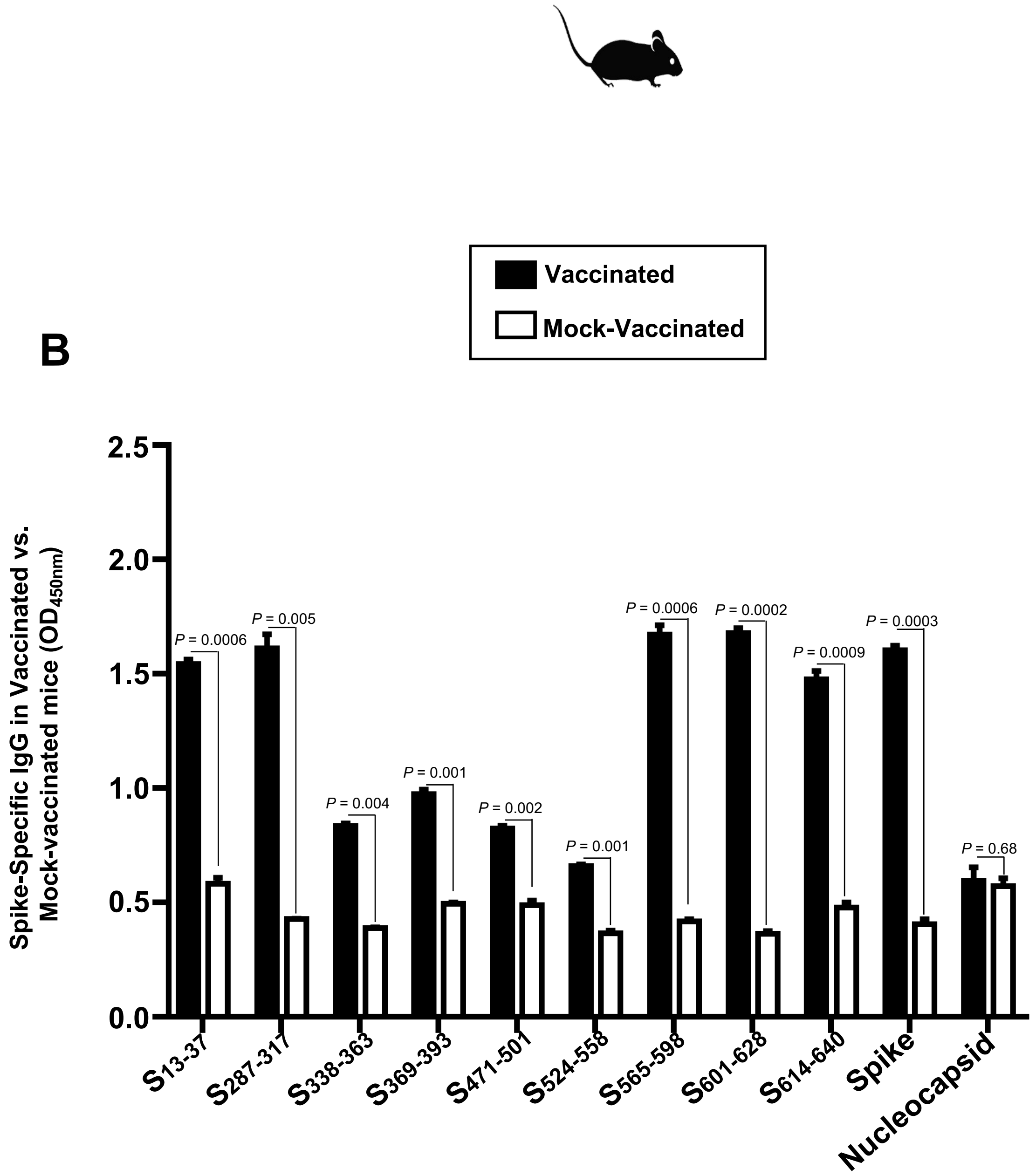
B
Vaccinated
Mock-Vaccinated
Spike-Specific IgG in Vaccinated vs. Mock-vaccinated mice (OD450nm)
P = 0.0006
P = 0.005
P = 0.004
P = 0.001
P = 0.002
P = 0.001
P = 0.0006
P = 0.0002
P = 0.0009
P = 0.0003
P = 0.68
S13-37
S287-317
S338-363
S369-393
S471-501
S524-558
S565-598
S601-628
S614-640
Spike
Nucleocapsid
